# Elevational variation in gut microbial community composition and predicted functional potential across three Himalayan redstart species

**DOI:** 10.64898/2026.11.09.765477

**Authors:** Pritam Chhetri, Nishma Dahal, Anil Kumar, Suruchi Rai, Surajit Bhattacharjee, Rakshak Kumar, Sangeet Lamichhaney

**Author notes:** Contributed equally (Pritam Chhetri, Nishma Dahal).

## Abstract

Environmental gradients and seasonal migration may influence host-associated microbial communities, yet their contributions to gut microbiome variation in montane birds remain poorly understood. We investigated gut microbial communities across 1,300–4,200 m in three Himalayan redstarts *(Phoenicurus fuliginosus, P. leucocephalus, and P. frontalis)* exhibiting contrasting seasonal movement strategies. Using 16S rRNA gene sequencing of 46 fecal samples, we characterized microbial diversity, community composition, distance–decay relationships, co-occurrence networks, and predicted functional potential. Elevation was significantly associated with microbial community composition, explaining approximately 3% of variation, whereas host species showed no significant association after accounting for elevation. *P. fuliginosus* exhibited significant compositional differences among most sampled elevations, whereas *P. leucocephalus* showed significant positive distance–decay relationships. Alpha diversity showed no consistent elevational trend, while core microbial taxa and co-occurrence networks varied among elevational groups. PICRUSt2 analyses identified 28 predicted KEGG Level 3 pathways differing significantly among elevations, including pathways associated with amino acid, lipid, glycan, carbohydrate, and bile acid metabolism. Together, these findings highlight elevation as a significant correlate of gut microbial community composition and predicted functional potential, with distinct patterns of microbial turnover among closely related Himalayan redstarts.

## Introduction

The gut microbiome represents a complex host-associated microbial community that contributes to nutrient metabolism, immune regulation, and physiological homeostasis (Rooks and Garrett, 2016; Smits *et al*., 2017). The diversity and composition of these communities are shaped by interactions among host evolutionary history, dietary ecology, physiological characteristics, and environmental conditions (Youngblut *et al*., 2019; Song *et al*., 2020). Although host-associated selective processes may promote the persistence of particular microbial lineages, environmental filtering, microbial dispersal, resource availability, and ecological interactions also contribute to gut microbial community assembly and turnover (Pereira and Berry, 2017). These processes can influence not only taxonomic composition but also the distribution of microbial functional traits. Understanding how environmental heterogeneity interacts with host-associated factors to shape microbial communities is therefore an important question in microbial ecology, particularly in natural populations occupying environmentally diverse landscapes.

Elevational gradients provide natural settings for investigating how environmental heterogeneity influences host-associated microbial communities (Pinos *et al*., 2025). Increasing elevation is accompanied by declining temperature and atmospheric oxygen partial pressure, together with changes in vegetation, habitat structure, and food resources (Li *et al*., 2026). These environmental differences may influence gut microbial communities directly through variation in environmental microbial exposure or indirectly through changes in host diet, physiology, and ecological interactions (Peng *et al*., 2025). However, studies examining gut microbial diversity across elevational gradients have reported contrasting patterns, with some documenting increased alpha diversity at higher elevations (Li *et al*., 2019), while others have reported declining diversity, including in human populations (Peng *et al*., 2025). Studies of high-elevation herbivores, including Tibetan pigs, Tibetan donkeys, and plateau pikas, have also suggested associations between gut microbial composition, dietary specialization, and metabolic functions related to nutrient utilization (Zhang *et al*., 2016; Li *et al*., 2019; Guo *et al*., 2022; Zhao *et al*., 2023). Together, these findings indicate that elevational patterns of gut microbial diversity are not universal but may depend on interactions among environmental conditions, host evolutionary history, dietary ecology, and exposure history (Youngblut *et al*., 2019; Yin *et al*., 2026).

Understanding elevational patterns in gut microbial communities is further complicated by differences in sampling design and host ecological characteristics. Many previous studies have compared populations occupying discrete low- and high-elevation environments (Li and Zhao, 2015; Wu *et al*., 2020), whereas studies incorporating multiple sampling locations across elevational gradients have revealed more complex and potentially species-specific patterns of microbial diversity and community composition (Li *et al*., 2019; Suzuki, Martins, *et al*., 2019). These differences suggest that comparisons between elevational extremes alone may not fully capture patterns of microbial community turnover across heterogeneous mountain landscapes. Furthermore, changes in overall microbial community composition may arise from shifts in multiple bacterial taxa rather than consistent changes in a few dominant microbial lineages. Examining community turnover across elevations can therefore provide insights into how microbial assemblages vary along environmental gradients and whether these patterns differ among host species. Despite growing interest in elevational microbiome ecology, studies incorporating multiple elevations and comparing microbial community responses among closely related wild bird species remain relatively limited compared with those involving mammals (but see Zhao *et al*., 2026).

Birds provide an informative system for examining how environmental heterogeneity and host ecology interact to shape gut microbial communities (Bodawatta *et al*., 2022). Avian gut microbiomes are influenced by dietary ecology, habitat use, phylogenetic relationships, and movement patterns, which may affect both exposure to environmental microorganisms and the conditions governing microbial community assembly. Comparative studies have suggested that host phylogenetic relationships explain less variation in gut microbial composition among birds than among mammals, potentially reflecting differences in host physiology and evolutionary history associated with powered flight (Kohl, 2012; Song *et al*., 2020; Bodawatta *et al*., 2022). Seasonal movements may further influence microbial communities by exposing birds to different environmental microbial pools, dietary resources, and habitat conditions, potentially contributing to microbial dispersal and community turnover (Skeen *et al*., 2023). Such processes may be particularly relevant in mountainous environments, where altitudinal migrants encounter changing environmental conditions and resource availability throughout their annual cycles (Ivy and Williamson, 2024).

Recent studies have documented seasonal and elevation-associated variation in avian gut microbial communities (Bo *et al*., 2025; Duan *et al*., 2026). However, whether closely related bird species occupying overlapping elevational ranges exhibit similar patterns of gut microbial community variation, despite differences in their habitat associations and movement ecology, remains poorly understood.

Beyond differences in microbial diversity and community composition, patterns of taxonomic overlap and functional potential provide complementary perspectives on the ecological organization of host-associated microbiomes. Identifying microbial taxa shared among host species and environmental conditions can reveal the extent of community overlap, whereas taxa restricted to particular hosts or environments may contribute to differences in microbial community composition. However, taxonomic turnover does not necessarily correspond to equivalent changes in microbial functional potential, as phylogenetically distinct microorganisms may possess overlapping metabolic capabilities (Louca *et al*., 2018). Conversely, shifts in microbial community composition may alter the representation of metabolic pathways involved in nutrient utilization, carbohydrate metabolism, amino acid metabolism, and other biochemical processes (Peng *et al*., 2025). Computational approaches enable the prediction of microbial gene families and metabolic pathways from 16S rRNA gene sequence data (Douglas *et al*., 2020), providing an indirect means of examining potential functional variation across ecological gradients. However, these predictions depend on reference genome representation and reflect inferred metabolic potential rather than directly measured microbial activity (Sun *et al*., 2020). Integrating taxonomic composition, patterns of shared microbial taxa, and predicted functional profiles may therefore provide a more comprehensive understanding of gut microbial variation across environmental gradients.

Himalayan redstarts (genus *Phoenicurus*) provide a useful comparative system for investigating gut microbial variation across environmentally heterogeneous montane landscapes. The Plumbeous Water Redstart *(Phoenicurus fuliginosus)* and White-capped Redstart *(P. leucocephalus)* are closely associated with streams and rivers, where they forage primarily in riparian habitats, whereas the Blue-fronted Redstart *(P. frontalis)* occupies a broader range of terrestrial montane habitats and is less dependent on flowing water (Buckton and Ormerod, 2008; Collar, 2020; Collar and Christie, 2020). These species also differ in their elevational distributions and seasonal movement patterns. *P. leucocephalus* and *P. frontalis* undertake seasonal elevational movements, whereas *P. fuliginosus* is generally resident or exhibits more localized elevational movements (IUCN, 2024). Their close evolutionary relationships, partially overlapping elevational distributions, and contrasting habitat associations provide an opportunity to examine how gut microbial communities vary among related host species occupying different ecological settings along an elevational gradient.

In this study, we characterized the gut microbial communities of three closely related Himalayan redstart species sampled across an elevational gradient spanning approximately 1,300–4,200 m in the eastern Himalayas. Using 16S rRNA gene amplicon sequencing and computational functional predictions, we investigated patterns of microbial diversity, community composition, taxonomic turnover, and predicted metabolic potential across elevations and among host species. Specifically, we aimed to (i) characterize variation in gut microbial alpha and beta diversity among elevations and host species; (ii) examine microbial community turnover and distance–decay relationships within species across elevational gradients; (iii) assess patterns of shared and unique microbial taxa among species and elevational groups; and (iv) evaluate variation in predicted metabolic pathways across elevations and host species. We hypothesized that differences in environmental conditions across elevations would be associated with variation in microbial community composition and predicted functional potential, with potentially contrasting patterns among host species reflecting differences in their ecological characteristics. We further predicted that some microbial taxa would be shared across species and elevations despite broader variation in community composition. By integrating taxonomic and predicted functional analyses, this study provides insights into patterns of gut microbial community organization across environmentally heterogeneous montane landscapes.

## Methods

### Study area and sampling

Field sampling was conducted during August and September 2025 across an elevational gradient in the West Kameng and Tawang districts of Arunachal Pradesh, eastern Himalayas, India. Six sampling locations spanning elevations from 1,300 to 4,200 m above sea level (a.s.l.) were selected to represent the observed elevational distributions of the target redstart species (*P. fuliginosus* and *P. leucocephalus*) during the post-breeding period, before the onset of seasonal downslope movements **(Fig. 1; Table S1).** Sampling was conducted following the southwest monsoon, when reduced stream and river water levels facilitated access to riparian habitats and mist-net deployment. This sampling period was also selected to reduce potential seasonal variation associated with breeding activities.

**Figure 1.**
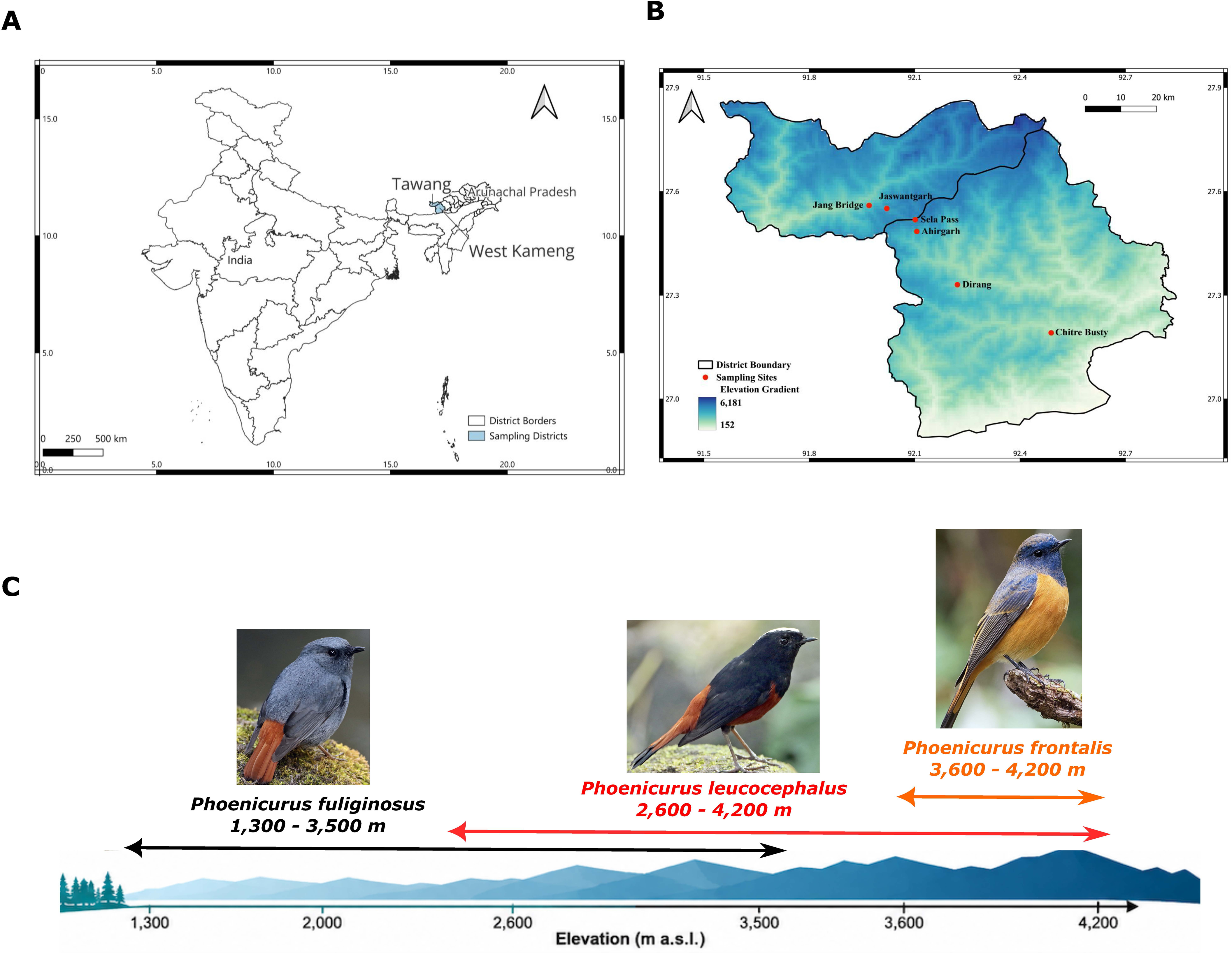
Geographic distribution of sampling sites and elevational ranges of three Himalayan redstart species in Arunachal Pradesh, India. (A) Map of India highlighting the study area in the West Kameng and Tawang districts of Arunachal Pradesh (shaded blue). Grey lines indicate district borders (B) Elevation map of the study area showing six sampling locations: Chitre Busty (1,300 m), Dirang (2,000 m), Jang Bridge (2,600 m), Jaswantgarh (3,500 m), Ahirgarh (3,600 m), and Sela Pass (4,200 m). Red circles indicate sampling locations, and background colors represent the elevational gradient. Black lines show district boundaries. (C) Schematic representation of the elevational ranges sampled for the three redstart species: Plumbeous water redstart (*Phoenicurus fuliginosus*, 1,300– 3,500 m), White-capped redstart (*P. leucocephalus*, 2,600–4,200 m), and Blue-fronted redstart (*P. frontalis*, 3,600–4,200 m). Bird images ©: Luke Seitz & Vincent Wang, eBird database. All elevations are expressed in meters above sea level (m a.s.l.).

Pre-sampling surveys confirmed the occurrence of *P. leucocephalus* between 2,600 and 4,200 m, with the lowest and highest observations recorded at Jang and Sela Pass, respectively. *P. fuliginosus* was observed between 1,300 and 3,800 m. However, mist-net deployment at the highest occupied elevation was not feasible because of steep and inaccessible terrain; consequently, the highest sampling location for this species was Jaswantgarh (3,500 m). Additional surveys below 1,300 m did not detect *P. fuliginosus*, suggesting that this elevation approximated the lower limit of its local distribution during the sampling period. Although *P. fuliginosus* is classified as resident in the IUCN Red List (IUCN, 2024), field observations suggested local elevational movements within the study area. We therefore describe this species as exhibiting local elevational movements rather than classifying it as a long-distance seasonal altitudinal migrant.

A total of 46 fresh faecal samples were collected from three redstart species: *P. fuliginosus* (n = 23), *P. leucocephalus* (n = 17), and *P. frontalis* (n = 6). Sampling primarily targeted *P. fuliginosus* and *P. leucocephalus* across their observed elevational distributions, whereas *P. frontalis* was encountered during field surveys and sampled opportunistically **(Table S1)**. Birds were captured using mist nets (12 × 2.5 m; 30-mm mesh) deployed along river corridors and other accessible habitats where individuals were observed actively foraging. Each captured bird was individually placed in a sterile paper bag for approximately 10 min to facilitate the collection of freshly voided faecal material. Following sample collection, standard morphometric measurements and body mass were recorded, and each individual was fitted with a uniquely numbered metal leg band before being released at its capture location. Fresh faecal samples were transferred into sterile 2-mL cryovials containing RNAlater (Sigma-Aldrich) for preservation of microbial nucleic acids. Samples were maintained in a portable cooler at approximately 4°C during fieldwork, transported to the laboratory on dry ice, and stored at −80°C until DNA extraction.

For comparative analyses, sampling locations were assigned to three elevation groups: low elevation (≤2,000 m), mid-elevation (2,600 m), and high elevation (≥3,500 m). These groups comprised 11, 11, and 24 samples, respectively, based on the site-level sample counts reported above. The elevation categories were used to facilitate comparisons of microbial diversity, community composition, and predicted functional potential among sampling environments.

### DNA extraction, 16S rRNA library preparation and sequencing

Total genomic DNA was extracted from 46 faecal samples using the QIAamp PowerFecal DNA Kit (Qiagen, Cat. No. 51604) following the manufacturer’s instructions. DNA concentration and purity were assessed by spectrophotometry, and DNA integrity was evaluated using agarose gel electrophoresis. The V3–V4 hypervariable region of the bacterial 16S rRNA gene was amplified using the universal bacterial primers S-D-Bact-0341-b-S-17 (5′-CCTACGGGNGGCWGCAG-3′) and S-D-Bact-0785-a-A-21 (5′-GACTACHVGGGTATCTAATCC-3′) (Klindworth *et al*., 2013). Amplicon libraries were prepared using a dual-indexing strategy with the Illumina Nextera CD Index Kit (8-bp indices). PCR products were purified using KAPA HyperPure Beads (Roche) to remove primer dimers and nonspecific amplification products. The resulting libraries, with an expected final fragment size of approximately 550 bp, were quantified, pooled at equimolar concentrations, and sequenced on an Illumina MiSeq platform (Eurofins Genomics) using 2 × 300-bp paired-end chemistry.

### Bioinformatics analysis

Raw sequencing reads were demultiplexed, and barcode sequences were removed before downstream processing. Initial sequence quality was assessed using FastQC v0.12.1 (https://www.bioinformatics.babraham.ac.uk/projects/fastqc/). Adapter sequences and low-quality bases (Phred quality score <30) were removed using Cutadapt v5.2(Martin, 2011), and the quality of the processed reads was reassessed before importing them into QIIME 2 v2026.1 (Bolyen *et al*., 2019).

Within QIIME 2, paired-end reads were processed using the DADA2 plugin (Callahan et al., 2016). Based on sequence quality profiles, forward reads were trimmed by 18 bp from the 5′ end and truncated at position 277, whereas reverse reads were truncated at position 200. DADA2 was used for quality filtering, sequencing error estimation and correction, paired-end read merging, and chimera detection and removal using the consensus method. The resulting high-resolution amplicon sequence variants (ASVs) were used to construct a feature table containing ASV abundances across samples.

Taxonomic classification of representative ASV sequences was performed using a naïve Bayes classifier trained on the SILVA 138.1 reference database (Quast *et al*., 2013) within QIIME2. The resulting ASV feature table and taxonomic assignments were used for subsequent analyses of microbial diversity, community composition, and differential taxon abundance. Sequencing depth and sampling completeness were evaluated using rarefaction analysis in QIIME 2. Rarefaction curves were generated by progressively subsampling reads from individual samples and examining changes in observed ASV richness with increasing sequencing depth. The extent to which curves approached an asymptote was used to assess whether sequencing depth adequately captured the detectable microbial diversity within each sample (Lundin *et al*., 2012).

### Statistical analysis

#### Alpha diversity

Alpha diversity was calculated from the rarefied ASV feature table to standardize sequencing depth across samples. Six complementary metrics were used to characterize different aspects of microbial diversity: Observed ASV richness, Chao1 richness, the abundance-based coverage estimator (ACE), Shannon diversity, Simpson diversity, and Faith’s phylogenetic diversity (PD). Observed ASV richness represents the number of distinct amplicon sequence variants (ASVs) detected within each sample, whereas Chao1 and ACE estimate richness by accounting for the contribution of rare ASVs (Chao, 1984; Chao and Lee, 1992). Shannon diversity incorporates both richness and evenness, while Simpson diversity places greater emphasis on dominant ASVs (Simpson, 1949; Spellerberg and Fedor, 2003). Faith’s PD incorporates evolutionary relationships among ASVs by measuring the total phylogenetic branch length represented within each sample (Faith, 1992).

#### Beta diversity and community composition

Beta diversity was assessed using Bray–Curtis and binary Jaccard dissimilarity matrices (Bray and Curtis, 1957) calculated from the rarefied ASV feature table. Bray–Curtis dissimilarity quantifies differences in ASV abundances between samples, whereas Jaccard dissimilarity evaluates differences in ASV presence and absence (Jaccard, 1901). Together, these metrics provide complementary assessments of abundance-based community variation and differences in microbial community membership. Patterns of microbial community composition were visualized using distance-based redundancy analysis (dbRDA) based on Bray– Curtis and Jaccard dissimilarities.

Associations between microbial community composition, elevation, and host species were evaluated using marginal permutational multivariate analysis of variance (PERMANOVA), implemented with the adonis2 function in the R package vegan (Oksanen *et al*., 2022). Separate models were fitted using Bray–Curtis and binary Jaccard dissimilarity matrices, with elevation treated as a continuous predictor and host species as a categorical predictor *(microbial distance ∼ elevation + host species)*. Marginal effects were evaluated using 999 permutations, allowing the association of each predictor with microbial community composition to be assessed while accounting for the other predictor. The proportion of variation explained by each predictor was summarized using R² values. Homogeneity of multivariate dispersion was assessed using permutational analysis of multivariate dispersions (PERMDISP) (Anderson, 2006), with significance evaluated using 999 permutations. Because host species were unevenly distributed across sampling elevations and each elevation corresponded to a distinct sampling location, the marginal PERMANOVA results were interpreted as conditional associations rather than independent causal effects of elevation or host species.

To further investigate elevational variation within host species, pairwise PERMANOVA analyses were conducted among sampled elevations separately for *P. fuliginosus* and *P. leucocephalus*. Corresponding PERMDISP analyses were performed to evaluate whether differences in multivariate dispersion might contribute to observed patterns of community differentiation. The multiple response permutation procedure (MRPP) was additionally used as a complementary nonparametric approach to assess differences in microbial community composition among groups (Mielke and Berry, 2001). P values from pairwise comparisons were adjusted using the Benjamini–Hochberg false discovery rate procedure, and adjusted P values < 0.05 were considered statistically significant.

#### Elevation-associated distance–decay analysis of gut microbial communities

To investigate whether gut microbial communities became increasingly dissimilar with greater elevational separation, we conducted distance–decay analyses separately for *P. fuliginosus* and *P. leucocephalus*, following (Li *et al*., 2019). *P. frontalis* was excluded because its samples were collected across a relatively narrow elevational range, limiting the ability to evaluate elevation-associated patterns. Pairwise microbial community dissimilarities were calculated from rarefied ASV counts using Bray– Curtis and binary Jaccard indices, representing abundance-based and presence–absence-based differences, respectively. Pairwise elevational distances were calculated as the absolute differences in sampling elevation (m above sea level) between individuals.

Associations between microbial community dissimilarity and elevational separation were evaluated using Mantel tests based on Spearman rank correlations with 9,999 permutations. Positive Mantel correlation coefficients indicate increasing microbial community dissimilarity with greater elevational separation, consistent with an elevation-associated distance–decay relationship. For visualization, pairwise elevational distances were plotted against the corresponding Bray–Curtis and Jaccard dissimilarities, with fitted linear regression lines and 95% confidence intervals used to illustrate overall trends. Because pairwise observations are not statistically independent, these regression lines and confidence intervals were interpreted descriptively, whereas statistical inference was based on permutation-derived Mantel test results. Mantel correlation coefficients (r) and permutation-based P values were used to summarize the strength and statistical significance of the associations.

Because sampling elevations corresponded to distinct geographic locations, elevational separation was inherently associated with differences in sampling site and potentially other environmental characteristics. Accordingly, distance–decay relationships were interpreted as associations between microbial community dissimilarity and elevational separation rather than evidence of an independent causal effect of elevation.

#### Taxonomic composition and genus-level associations with elevation

Microbial taxonomic composition was characterized by calculating the relative abundances of bacterial phyla and genera, expressed as the proportion of reads assigned to each taxon relative to the total number of taxonomically classified reads within each sample. Taxonomic profiles were summarized across host species and elevational groups to characterize broad patterns of bacterial community composition. For descriptive comparisons, relative abundances were first averaged across individual samples within each species–elevation sampling group. These group-level values were subsequently summarized separately by host species and elevation category (low, mid, and high). For each taxon, the arithmetic mean and standard deviation were calculated across the corresponding group-level relative abundances, giving equal weight to each contributing species–elevation group rather than to individual samples. Because the number of contributing groups was limited, these summaries were used to describe broad compositional patterns rather than for formal statistical inference.

To investigate associations between bacterial genera and elevational variation, genus-level relative abundances were analyzed separately for *P. fuliginosus and P. leucocephalus*. Genera with a mean relative abundance ≥0.1% across samples within the respective host species were retained for analysis, following (Suzuki, Martins, *et al*., 2019). This filtering threshold was applied independently to each species to account for differences in microbial community composition. Associations between genus-level relative abundance and sampling elevation were evaluated using Spearman rank correlations. One *P. leucocephalus* sample was excluded from the genus-level correlation analysis. To account for multiple comparisons, P-values were adjusted separately within each host species using the Benjamini–Hochberg false discovery rate procedure (Benjamini and Hochberg, 1995). Adjusted P-values < 0.05 were considered statistically significant. Statistical analyses and visualizations were performed in R v4.5.3, with graphical representations generated using the ggplot2 package (Wickham, 2016).

#### Core microbiome analysis

Core microbiome analyses were conducted at the phylum and genus levels to identify bacterial taxa consistently represented within host species and across elevational groups. Taxonomic assignments were based on the SILVA 138.1 reference database. ASVs classified as chloroplasts or mitochondria were excluded before taxonomic aggregation. Taxonomic annotations lacking informative assignments at a given rank (e.g., “uncultured”, “unidentified”, “metagenome”, or “unclassified”) were treated as unresolved at that rank and retained at the lowest confidently assigned higher taxonomic level. ASV counts were aggregated separately at the phylum and genus levels and converted to relative abundances within each sample.

Core taxa were defined using two criteria: occurrence in at least 70% of samples within a group and a mean relative abundance of at least 0.1% across samples in that group. Core taxa were identified independently for each host species and for three predefined elevational categories: low (≤2,000 m), mid (2,600 m), and high (≥3,500 m). The number and taxonomic identity of core taxa were compared among host species and elevational categories. Taxa shared across all three elevational categories were identified through intersections of their respective core taxon sets, and overlap among host species and elevational categories was visualized using UpSet plots. These comparisons were descriptive and were not used to infer statistically significant differences in core community composition.

For genus-level comparisons, genera identified as core in at least one host species or elevational category were retained, and their relative abundances were summarized for each host species within each elevational category. Species–elevation groups containing fewer than two samples were excluded from these summaries. Relative abundances were log-transformed after adding a pseudocount of 0.0001, and hierarchical clustering was applied to both genera and species–elevation groups to visualize similarities in their relative-abundance profiles. Clustering patterns were interpreted descriptively rather than as formal evidence of ecological differentiation among host species or elevations.

#### Functional prediction and analysis

The functional potential of gut microbial communities was inferred from the unrarefied ASV abundance table using PICRUSt2 (Douglas *et al*., 2020). Hidden-state prediction was performed using maximum parsimony, with a maximum Nearest Sequenced Taxon Index (NSTI) threshold of 2. Predicted abundances of KEGG Orthologs (KOs) were generated and converted to sample-wise relative abundances in R. KO annotations were mapped to the Kyoto Encyclopedia of Genes and Genomes (KEGG) functional hierarchy, and predicted functions were summarized at KEGG Level 3 to characterize variation in inferred microbial functional potential across elevational groups.

Differences in predicted KEGG Level 3 functional abundances among the three predefined elevation categories (low, ≤2,000 m; mid, 2,600 m; and high, ≥3,500 m) were evaluated using Kruskal–Wallis tests, followed by pairwise Wilcoxon rank-sum tests. Associations between predicted functional abundances and sampling elevation were further evaluated using Spearman rank correlations to identify monotonic elevational trends. To account for multiple comparisons, P-values were adjusted using the Benjamini–Hochberg false discovery rate procedure, with adjusted P-values (q) < 0.05 considered statistically significant.

Because functional profiles were inferred from 16S rRNA gene sequences rather than directly measured through shotgun metagenomics or meta-transcriptomics, the results were interpreted as predictions of microbial functional potential rather than evidence of actual metabolic activity. Data processing, statistical analyses, and visualization were performed in R v4.5.3 using the *tidyverse* collection of packages, including *dplyr, tidyr, readr, stringr, purrr, tibble, and ggplot2* (Wickham *et al*., 2019).

#### Co-occurrence network construction and putative keystone-taxon identification

To investigate patterns of co-occurrence among gut bacterial taxa and identify potentially influential members of microbial communities across the elevational gradient, genus-level association networks were constructed using relative-abundance data. Networks were generated for the complete dataset and separately for each of the three predefined elevation categories (low, ≤2,000 m; mid, 2,600 m; and high, ≥3,500 m). Elevation-specific networks were examined to characterize variation in microbial association patterns among elevational zones. This approach was motivated by observed elevation-associated differences in microbial community composition, whereas host-species effects were not statistically significant in the preceding community-level analyses. Networks were not constructed separately for individual host species because of the limited and uneven sample sizes.

To reduce the influence of sparsely represented taxa, genera were retained if they occurred in at least 20% of samples within the respective dataset. This threshold corresponded to a minimum prevalence of 10 samples for the complete dataset (n = 46), three samples each for the low- and mid-elevation groups (n = 11 per group), and five samples for the high-elevation group (n = 24). Pairwise associations among retained genera were estimated using Spearman rank correlations, and P-values were adjusted for multiple comparisons using the Benjamini–Hochberg false discovery rate procedure. Associations with an absolute Spearman correlation coefficient |ρ| ≥ 0.70 and adjusted P < 0.05 were retained as network edges (Bauchinger *et al*., 2024). Positive and negative correlations represented taxa exhibiting concordant and opposing relative-abundance patterns, respectively.

Networks were constructed and analyzed using the *igraph* package in R. Network topology was characterized using degree, betweenness centrality, closeness centrality, eigenvector centrality, and node strength, calculated for the largest connected component of each network. Genera with both degree and betweenness centrality values at or above the 90th percentile within their respective networks were classified as putative keystone taxa. Network visualization was performed using Gephi (El-Badry *et al*., 2026).

Because network edges were inferred from correlations among compositional relative-abundance data, they were interpreted as statistical associations rather than direct evidence of ecological interactions. Likewise, putative keystone taxa were identified based on their topological prominence within the inferred networks rather than experimentally demonstrated functional importance. Given the relatively small and unequal sample sizes across elevation groups, differences in network structure and keystone-taxon identity were interpreted descriptively and not as definitive evidence of elevational changes in microbial interactions.

## Results

### Gut microbiome diversity across elevation bands

Following quality filtering, denoising, paired-end read merging, and chimera removal, 581,417 high-quality reads were retained from 46 fecal samples, with 4,705–22,772 non-chimeric reads per sample **(Table S2)**. Rarefaction curves approached saturation with increasing sequencing depth, with most showing substantial flattening at approximately 10,000 sequences **(Fig. S1)**, suggesting that sequencing captured a substantial proportion of the detectable bacterial diversity.

Alpha diversity differed significantly among the nine species–elevation groups across all six metrics (Kruskal–Wallis P = 0.0167 for observed ASVs and Chao1, 0.0168 for ACE, 0.0174 for Simpson, 0.0223 for Faith’s phylogenetic diversity [PD], and 0.0240 for Shannon; BH-adjusted P = 0.024 for all metrics; **Fig. 2**). Across metrics, *P. fuliginosus* at 2,600 m generally exhibited the lowest diversity, whereas *P. leucocephalus* and *P. frontalis* at 4,200 m exhibited the highest median richness and diversity. Dunn’s post hoc comparisons indicated lower diversity in *P. fuliginosus* at 2,600 m than in both species at 4,200 m for observed ASVs, Chao1, ACE, Shannon, and Simpson diversity, whereas Faith’s PD differed only between *P. fuliginosus* at 2,600 m and *P. leucocephalus* at 4,200 m. Other species–elevation groups generally showed intermediate values without statistically distinguishable differences from either extreme. Within *P. fuliginosus*, diversity declined from 1,300–2,000 m to a minimum at 2,600 m before increasing at 3,500 m, although substantial variation was observed among individuals. Simpson diversity remained relatively high (approximately 0.9–1.0) across most groups.

**Figure 2.**
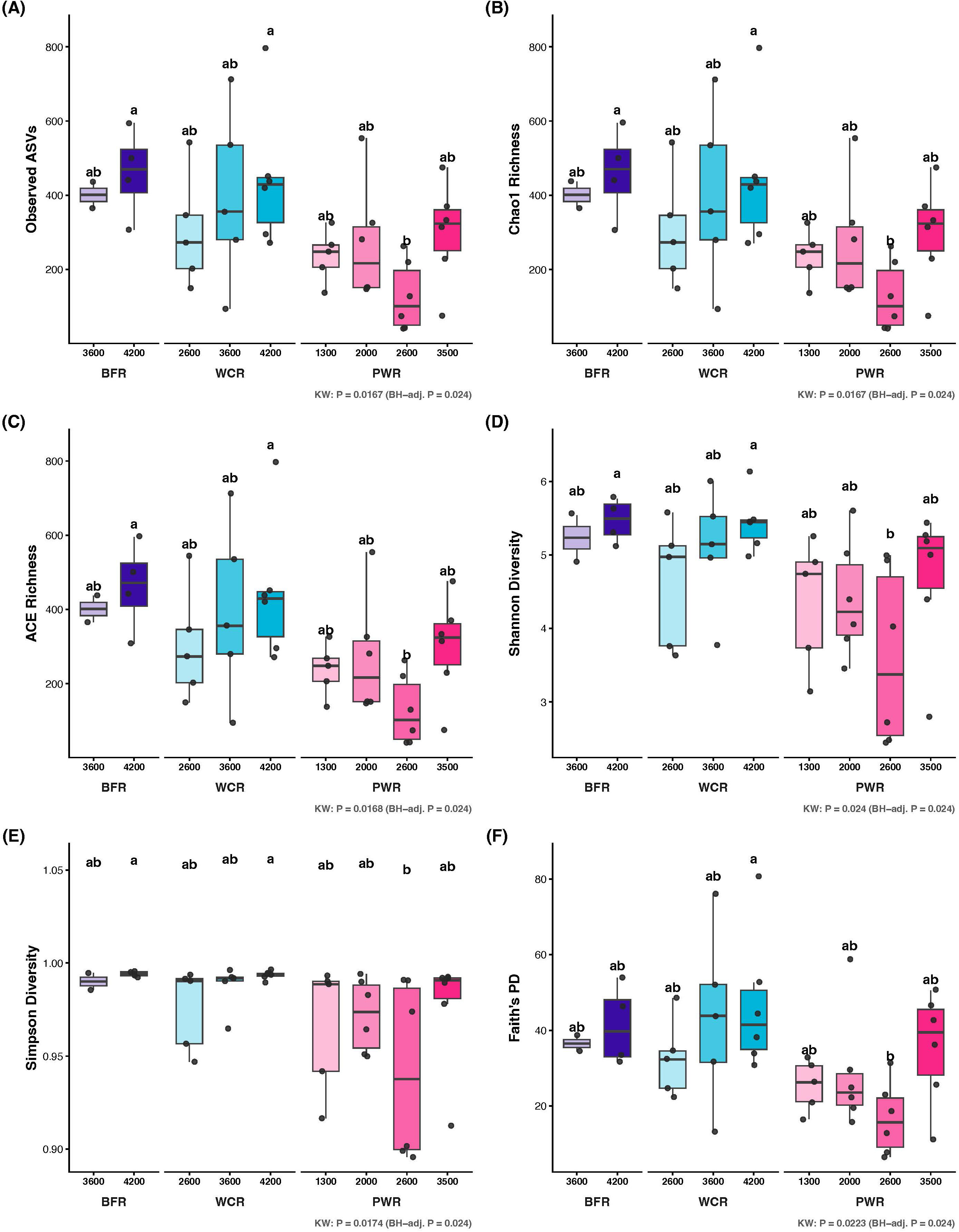
Gut microbial alpha diversity across host species and sampling elevations. Alpha diversity was assessed using six metrics: **(A)** observed Amplicon Sequence Variants (ASVs), **(B)** Chao1 richness, **(C)** ACE richness, **(D)** Shannon diversity, **(E)** Simpson diversity, and **(F)** Faith’s phylogenetic diversity (PD). Samples are grouped by host species and sampling elevation (m a.s.l.). Host species are abbreviated as BFR, Blue-fronted redstart (*Phoenicurus frontalis*); WCR, White-capped redstart (*P. leucocephalus*); and PWR, Plumbeous water redstart (*P. fuliginosus*). Boxes represent the interquartile range (IQR), with horizontal lines indicating medians; whiskers extend to 1.5 × IQR, and individual points represent samples from individual birds. The Kruskal–Wallis test P value and corresponding Benjamini–Hochberg (BH)-adjusted P value are shown below each panel. Letters indicate groupings based on Dunn’s post hoc pairwise comparisons with BH correction; groups sharing at least one letter are not significantly different (adjusted P ≥ 0.05).

Within-species comparisons across elevations revealed limited evidence of differences in alpha diversity **(Data Table 1)**. In *P. fuliginosus*, diversity at 2,600 m was lower than at 3,500 m for observed ASVs, Chao1, and ACE (Dunn’s Z = −2.24, P = 0.025), Shannon diversity (Z = −2.08, P = 0.037), and Faith’s PD (Z = −2.08, P = 0.037), whereas Simpson diversity did not differ (P = 0.078). Richness at 2,600 m also tended to be lower than at 1,300 m (P = 0.068) and 2,000 m (P = 0.055). In *P. leucocephalus*, only Simpson diversity differed between 2,600 and 4,200 m (Z = −2.01, P = 0.045).

**Table 1.** Pairwise comparisons of gut microbial beta diversity among elevations within *P. fuliginosus* (1300 m, 2000 m, 2600 m, and 3500 m) and *P. leucocephalus* (2600 m, 3600 m, and 4200 m) based on Bray-Curtis and Jaccard dissimilarities. Community differences were evaluated using PERMANOVA, while homogeneity of multivariate dispersion was assessed using PERMDISP. Significant results are indicated by bold adjusted P values.

| *Phoenicurus fuliginosus* - **Bray Curtis** |  |  |  |  |  |
|---|---|---|---|---|---|
| **Elevation Group** |  | **PERMANOVA** |  | **PERMDISP** |  |
| **Group1** | **Group2** | **R<sup>2</sup>** | **P** | **F** | **P** |
| **1300** | 2000 | 0.167713 | **0.009** | 0.297 | 0.915 |
| **1300** | 2600 | 0.14975 | **0.0168** | 0.623585 | 0.915 |
| **1300** | 3500 | 0.143203 | **0.009** | 2.474874 | 0.816 |
| **2000** | 2600 | 0.136058 | **0.012** | 0.135688 | 0.918 |
| **2000** | 3500 | 0.128802 | **0.012** | 0.418011 | 0.915 |
| **2600** | 3500 | 0.111373 | 0.165 | 0.001844 | 0.952 |
| *Phoenicurus fuliginosus* - **Jaccard** |  |  |  |  |  |
| **Elevation Group** |  | **PERMANOVA** |  | **PERMDISP** |  |
| **Group1** | Group2 | R2 | P | F | P |
| **1300** | 2000 | 0.136 | **0.008** | 5.529011 | 0.064 |
| **1300** | 2600 | 0.127 | **0.008** | 8.489873 | 0.064 |
| **1300** | 3500 | 0.127 | **0.008** | 7.519062 | 0.064 |
| **2000** | 2600 | 0.103 | **0.0204** | 1.13344 | 0.489 |
| **2000** | 3500 | 0.108 | **0.009** | 0.310579 | 0.595 |
| **2600** | 3500 | 0.097 | 0.177 | 0.369899 | 0.595 |
| *Phoenicurus Leucocephalus* - **Bray Curtis** |  |  |  |  |  |
| **Elevation Group** |  | **PERMANOVA** |  | **PERMDISP** |  |
| **Group1** | Group2 | R2 | P | F | P |
| **2600** | 3600 | 0.123 | 0.176 | 0.693 | 0.51 |
| **2600** | 4200 | 0.123 | 0.096 | 0.0006 | 0.989 |
| **3600** | 4200 | 0.112 | 0.156 | 0.951 | 0.514 |
| *Phoenicurus Leucocephalus* - **Jaccard** |  |  |  |  |  |
| **Elevation Group** |  | **PERMANOVA** |  | **PERMDISP** |  |
| **Group1** | Group2 | R2 | P | F | P |
| **2600** | 3600 | 0.114 | 0.2 | 0.547 | 0.481 |
| **2600** | 4200 | 0.109 | 0.06 | 1.29 | 0.346 |
| **3600** | 4200 | 0.105 | 0.147 | 4.03 | 0.097 |

However, none of these within-species pairwise differences remained statistically significant after BH correction across all diversity metrics (adjusted P ≥ 0.22 for *P. fuliginosus* and ≥ 0.52 for *P. leucocephalus*). Elevational comparisons were not performed for *P. frontalis* because of its limited representation at 3,600 m (n = 2) and 4,200 m (n = 4).

Comparisons among host species within shared elevational bands also revealed few differences **(Data Table 1)**. At 2,600 m, *P. fuliginosus* (n = 6) exhibited lower observed ASV richness, Chao1, and ACE than P. leucocephalus (n = 5; χ² = 4.03, Z = −2.01, P = 0.045), as well as lower Faith’s PD (χ² = 4.80, Z = −2.19, P = 0.028). Shannon and Simpson diversity did not differ significantly between species (P = 0.144 and 0.201, respectively). No significant differences were detected among the three host species at 3,500–3,600 m or between *P. frontalis* and *P. leucocephalus* at 4,200 m. After BH correction across all diversity metrics, the nominal differences observed at 2,600 m were no longer significant (adjusted P = 0.335), and no significant differences were detected at the higher elevational bands (adjusted P ≥ 0.81).

### Gut microbial community composition

Marginal PERMANOVA revealed significant associations between elevation and gut microbial community composition after accounting for host species. Elevation explained 3.33% of the variation in Bray–Curtis dissimilarity (R² = 0.0333, P = 0.003) and 2.86% of the variation in Jaccard dissimilarity (R² = 0.0286, P = 0.001). In contrast, host species was not significantly associated with microbial community composition after accounting for elevation, although it explained 4.26% of the variation in Bray–Curtis dissimilarity (R² = 0.0426, P = 0.585) and 4.47% of the variation in Jaccard dissimilarity (R² = 0.0447, P = 0.284).

Distance-based redundancy analysis (dbRDA) ordinations were broadly consistent with these findings **(Fig. 3)**. Samples exhibited partial separation along the first constrained ordination axis (dbRDA1), which accounted for 48.7% and 41.7% of the constrained variation in the Bray–Curtis and Jaccard analyses, respectively. The fitted elevation vector was closely aligned with this axis, indicating an association between elevational variation and microbial community composition. In contrast, samples did not form clearly distinguishable clusters according to host species.

**Figure 3.**
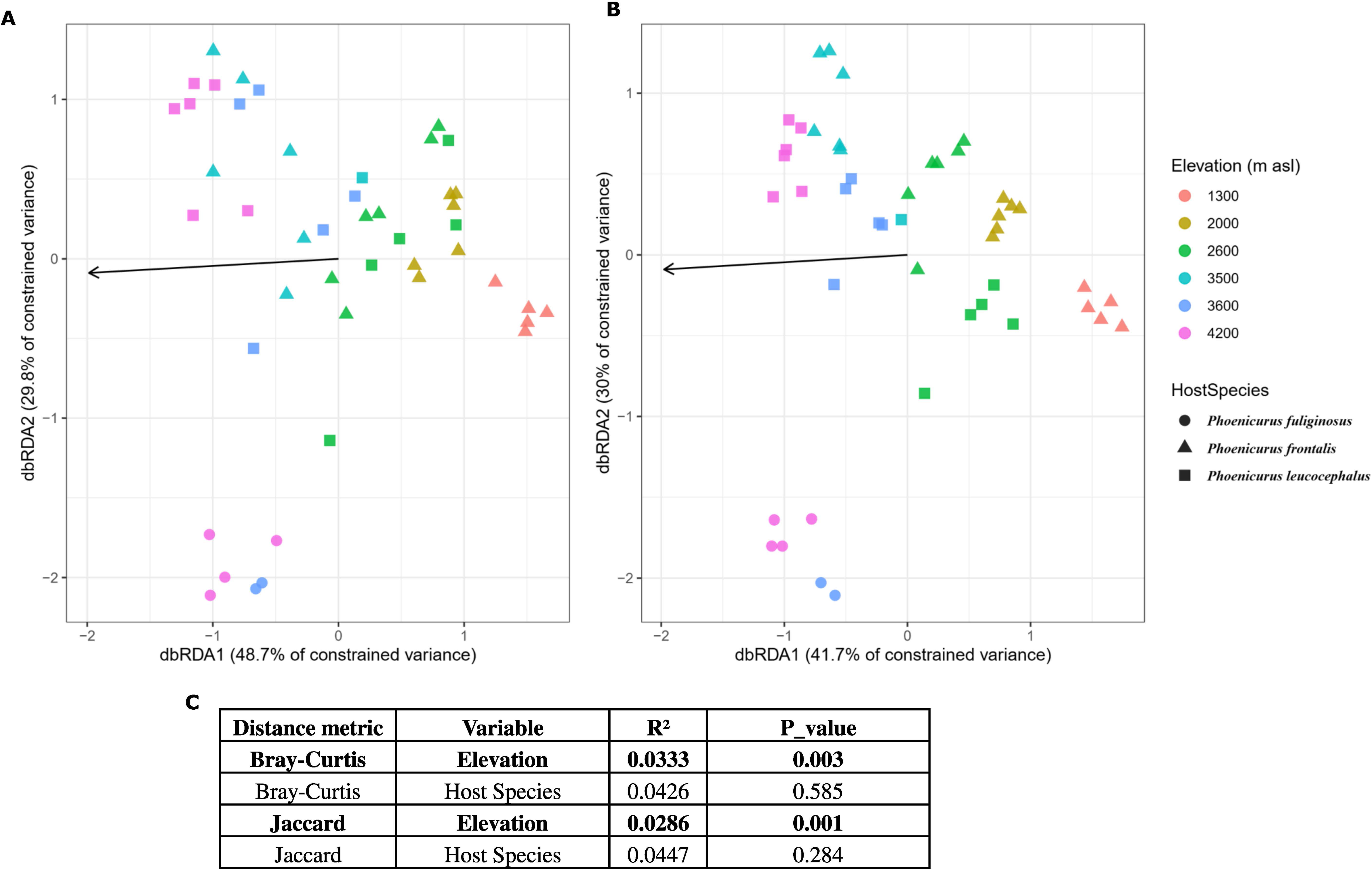
Variation in gut microbial community composition across sampling elevations and host species. Distance-based redundancy analysis (dbRDA) ordinations based on **(A)** Bray–Curtis and **(B)** Jaccard dissimilarities. Each point represents an individual bird, with colors indicating sampling elevation (m a.s.l.) and symbols indicating host species (circles, *Phoenicurus frontalis*; triangles, *P. fuliginosus*; squares, *P. leucocephalus*). Axis labels indicate the percentage of constrained variation explained by each dbRDA axis. Arrows represent the fitted elevation vector. **(C)** Marginal permutational multivariate analysis of variance (PERMANOVA; *adonis2*, 999 permutations) assessing the associations of elevation and host species with microbial community composition. The proportion of variation explained (R²) and corresponding P values are presented for each predictor, with statistically significant associations (P < 0.05) highlighted in bold.

### Pairwise comparisons of gut microbial community composition among host species within shared elevational bands

Pairwise PERMANOVA analyses were conducted to compare gut microbial community composition among host species sampled within shared elevational bands **(Table S3)**. At 2,600 m, no significant differences were detected between *P. fuliginosus* and *P. leucocephalus* using either Bray–Curtis (R² = 0.074, P = 0.982) or Jaccard dissimilarity (R² = 0.099, P = 0.502). PERMDISP tests also indicated no significant differences in multivariate dispersion between the two species (Bray–Curtis: P = 0.921; Jaccard: P = 0.226).

At 3,500–3,600 m, none of the pairwise comparisons among the three host species showed significant differences in microbial community composition (Bray–Curtis: R² = 0.077–0.155, P ≥ 0.550; Jaccard: R² = 0.087–0.151, P ≥ 0.364). However, PERMDISP revealed significant differences in multivariate dispersion in comparisons involving *P. frontalis*. For Bray–Curtis dissimilarity, significant dispersion differences were observed between *P. frontalis* at 3,600 m and *P. fuliginosus* at 3,500 m (F = 114.6, P = 0.0015), and between *P. frontalis* and *P. leucocephalus* at 3,600 m (F = 110.8, P = 0.0015). Corresponding comparisons using Jaccard dissimilarity were also significant (F = 304.1 and 462.2, respectively; both P = 0.0015). In contrast, dispersion did not differ significantly between *P. fuliginosus* at 3,500 m and *P. leucocephalus* at 3,600 m (Bray–Curtis: P = 0.974; Jaccard: P = 0.068).

At 4,200 m, the comparison between *P. frontalis* and *P. leucocephalus* approached statistical significance for Bray–Curtis dissimilarity (R² = 0.121, P = 0.052), whereas the Jaccard-based comparison was not significant (R² = 0.114, P = 0.116). PERMDISP detected no significant difference in dispersion using Bray–Curtis dissimilarity (P = 0.415), but a significant difference was observed using Jaccard dissimilarity (F = 8.554, P = 0.011).

### Elevation-associated differences in gut microbial community composition within host species

Pairwise PERMANOVA analyses were performed to examine differences in gut microbial community composition among sampled elevations within *P. fuliginosus* and *P. leucocephalus* **(Table 1)**. In *P. fuliginosus*, five of the six possible pairwise elevation comparisons revealed significant differences in microbial community composition based on both Bray–Curtis and Jaccard dissimilarities. The only non-significant comparison was between 2,600 and 3,500 m. PERMDISP tests were non-significant for all pairwise comparisons, indicating no statistically detectable differences in multivariate dispersion among the elevation groups. In contrast, none of the six pairwise elevation comparisons in *P. leucocephalus* showed significant differences in microbial community composition after multiple-testing correction (Bray–Curtis: adjusted P = 0.096–0.176; Jaccard: adjusted P = 0.060–0.200). The comparison between 2,600 and 4,200 m yielded the lowest adjusted P value under Jaccard dissimilarity (P = 0.060). PERMDISP tests were non-significant across all pairwise comparisons, with no statistically detectable differences in multivariate dispersion among elevations.

### Distance-decay patterns of gut microbial communities

Mantel tests were performed to examine associations between elevational separation and gut microbial community dissimilarity within *P. fuliginosus* and *P. leucocephalus* **(Fig. 4)**. In *P. fuliginosus*, Bray–Curtis dissimilarity exhibited a weak positive but non-significant correlation with elevational distance (Mantel r = 0.113, P = 0.091). Similarly, Jaccard dissimilarity showed a positive association that did not reach statistical significance (Mantel r = 0.133, P = 0.059). In contrast, *P. leucocephalus* exhibited significant positive correlations between elevational distance and microbial community dissimilarity for both Bray–Curtis (Mantel r = 0.266, P = 0.009) and Jaccard dissimilarity (Mantel r = 0.280, P = 0.004).

**Figure 4.**
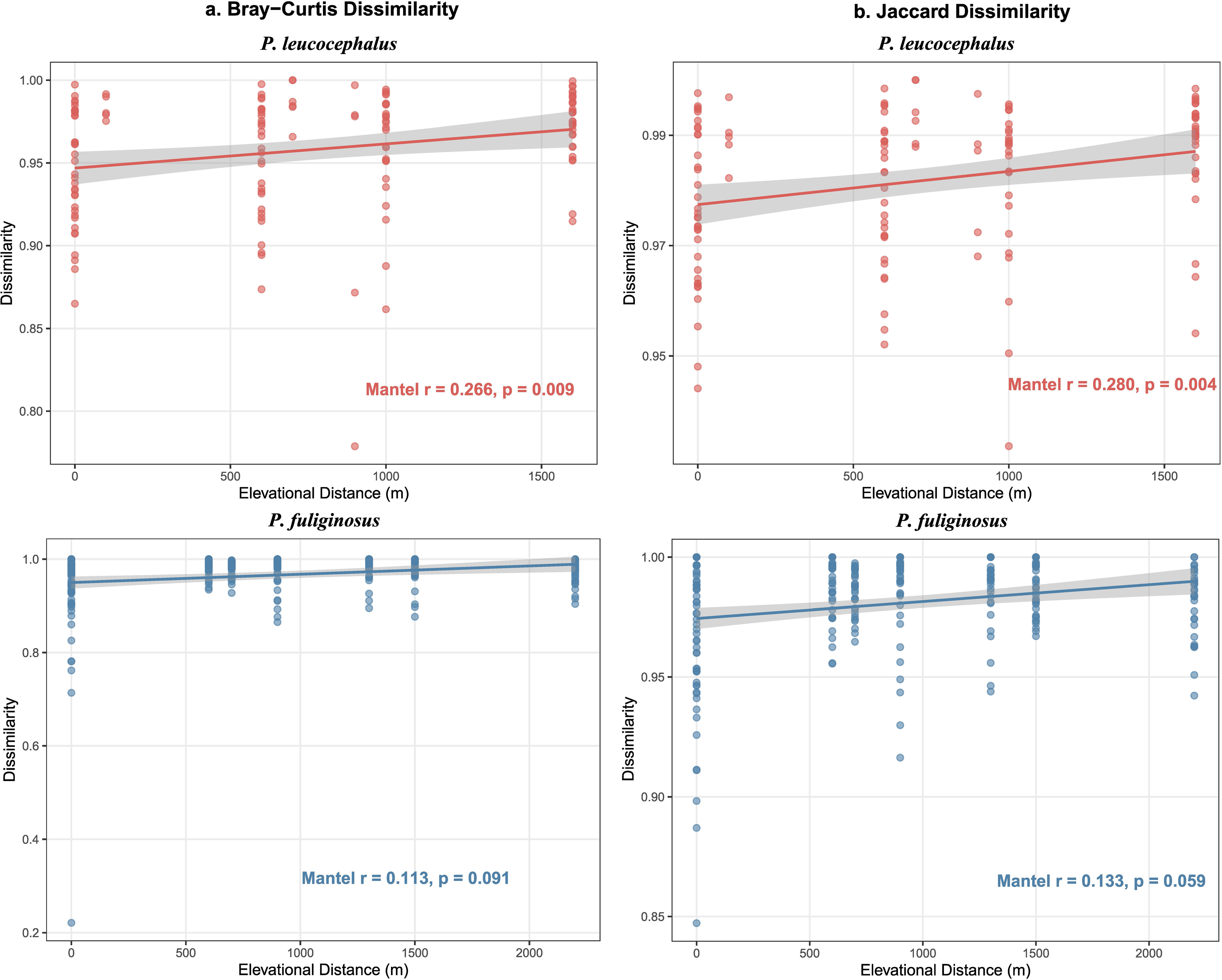
Distance–decay relationships between gut microbial community dissimilarity and elevational separation in two Himalayan redstart species. Relationships between pairwise elevational distance (m) and gut microbial community dissimilarity based on (a) Bray–Curtis and (b) Jaccard indices for White-capped redstart (*Phoenicurus leucocephalus*; upper panels, red) and Plumbeous water redstart (*P. fuliginosus*; lower panels, blue). Each point represents a pairwise comparison between individual birds. Solid lines indicate fitted linear regression trends, with shaded grey bands representing 95% confidence intervals. Mantel correlation coefficients (Spearman’s *r*) and corresponding P values, calculated using 9,999 permutations, are shown in each panel.

### Taxonomic composition and elevation-associated variation of the gut microbiome

At the phylum level, gut microbial communities across the three Himalayan redstart species were predominantly composed of Proteobacteria and Firmicutes, followed by Actinobacteriota and Bacteroidota, although their relative abundances varied among host species and elevational groups **(Fig. 5; Table S4)**. Proteobacteria was the most abundant phylum across most host species–elevation groups, with the highest mean relative abundance in *P. fuliginosus* (56.9% ± 11.5%), followed by *P. leucocephalus* (39.3% ± 15.4%) and *P. frontalis* (31.6% ± 8.8%). In contrast, Firmicutes showed the highest mean relative abundance in *P. frontalis* (34.2% ± 8.4%), followed by *P. leucocephalus* (31.1% ± 8.1%) and *P. fuliginosus* (21.4% ± 10.9%). Actinobacteriota exhibited a similar distribution, accounting for 15.1%, 14.3%, and 9.1% of the microbial communities in *P. frontalis*, *P. leucocephalus*, and *P. fuliginosus*, respectively. The relative abundances of less prevalent bacterial phyla are presented in **Table S4**. Across elevational categories, Proteobacteria reached its highest mean relative abundance at mid-elevation sites (64.4% ± 11.8%) and declined at high-elevation sites (34.8% ± 9.7%). Firmicutes exhibited the opposite pattern, reaching its highest mean relative abundance at high elevation (33.3% ± 5.7%; **Fig. 5; Table S4**).

**Figure 5.**
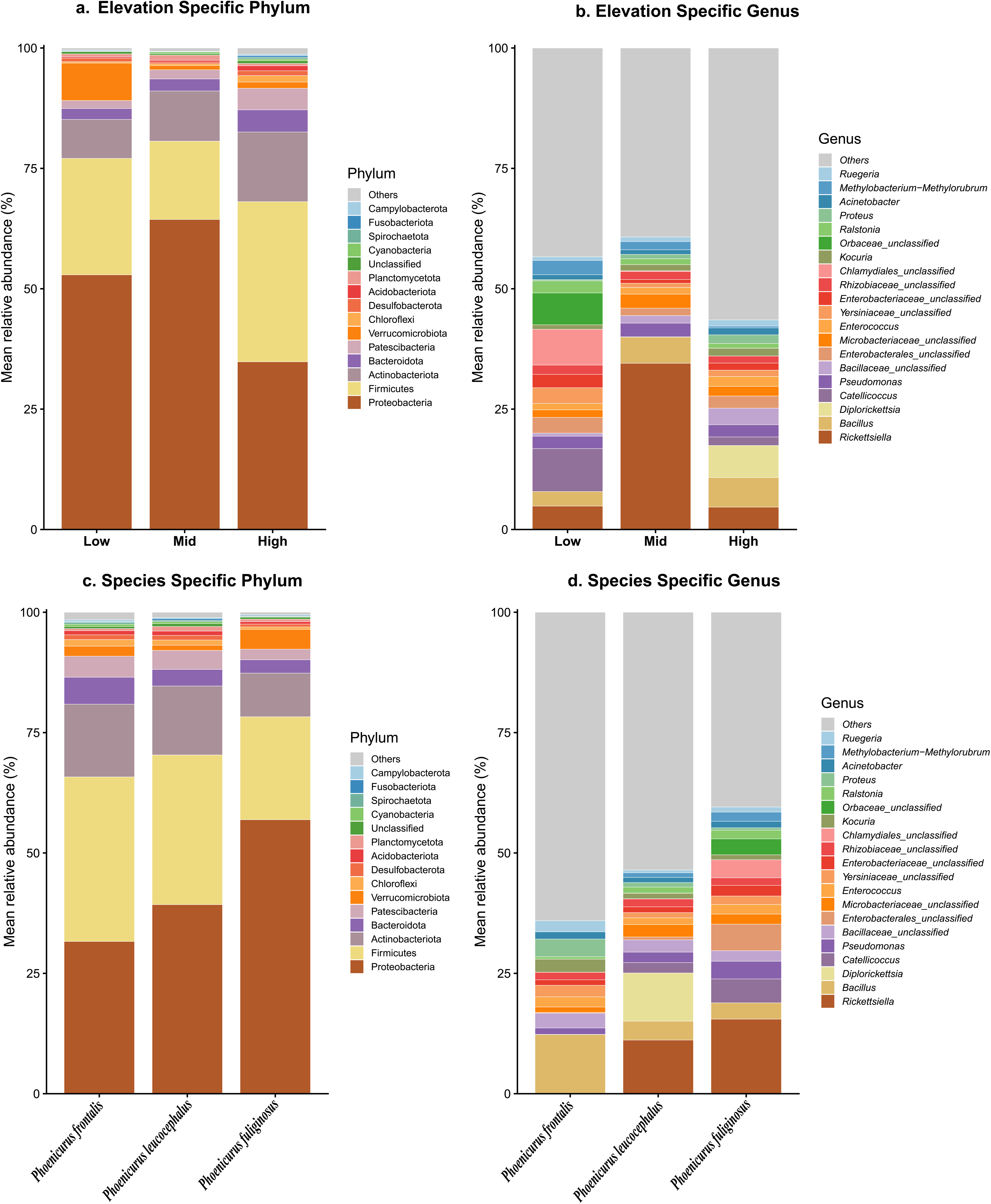
Taxonomic composition of gut microbial communities across elevational groups and host species in Himalayan redstarts. Mean relative abundances (%) of the dominant bacterial phyla (a, c) and genera (b, d), grouped by elevational category (a, b) and host species (c, d). Elevational categories were defined as low (≤2,000 m), mid (2,600 m), and high (≥3,500 m). Taxa with lower relative abundances were grouped as “Others”, whereas amplicon sequence variants (ASVs) that could not be assigned to the corresponding taxonomic rank were classified as “Unclassified”.

At the genus level, Rickettsiella varied considerably among host species, with mean relative abundances of 0.05% ± 0.03% in *P. frontalis*, 11.1% ± 12.9% in *P. leucocephalus*, and 15.5% ± 20.6% in *P. fuliginosus*. Across elevational categories, Rickettsiella was most abundant at mid-elevation sites (34.5% ± 16.0%), compared with low-(4.9% ± 6.7%) and high-elevation sites (4.6% ± 8.5%). Diplorickettsia was predominantly detected in *P. leucocephalus* (10.0% ± 20.0%) and was most abundant in the high-elevation group (6.7% ± 16.3%), with negligible relative abundances in the remaining groups. Spearman rank correlation analyses identified both positive and negative associations between the relative abundances of bacterial genera and elevation within *P. fuliginosus* and *P. leucocephalus* **(Fig. S3; Table S5)**. Following Benjamini–Hochberg correction for multiple testing, only Lactococcus exhibited a significant negative correlation with elevation in *P. fuliginosus* (adjusted P < 0.01). No genus showed a statistically significant association with elevation in *P. leucocephalus* after multiple-testing correction.

### Core microbiome variation across host species and elevation groups

Core microbiome analysis, based on a prevalence threshold of ≥70% and a mean relative abundance threshold of ≥0.1%, revealed variation in the number of core bacterial taxa among host species and elevational groups. At the host-species level, 9, 14, and 15 core phyla and 11, 44, and 36 core genera were identified in *P. fuliginosus, P. leucocephalus, and P. frontalis*, respectively. Across elevational groups, 9, 11, and 16 core phyla and 21, 16, and 43 core genera were identified at low, mid, and high elevations, respectively **(Fig. S4; Table S6)**.

UpSet analysis revealed that nine core phyla were shared among all three host species, and the same nine phyla were also shared across all three elevational groups **(Fig. 6**; **Table 2)**. At the genus level, eight core genera were shared among all three host species, whereas the overlap among elevational groups varied. The high-elevation group contained the largest number of unique core genera (29), followed by the low-(5) and mid-elevation groups (3). Among host species, *P. leucocephalus* had the highest number of unique core genera (20), followed by *P. frontalis* (12) and *P. fuliginosus* (1) **(Fig. 6; Table S6)**. Heatmap analysis further revealed differences in the relative abundances of core genera among host species and elevational groups, with several genera exhibiting contrasting abundance patterns across groups **(Fig. S4)**.

**Figure 6.**
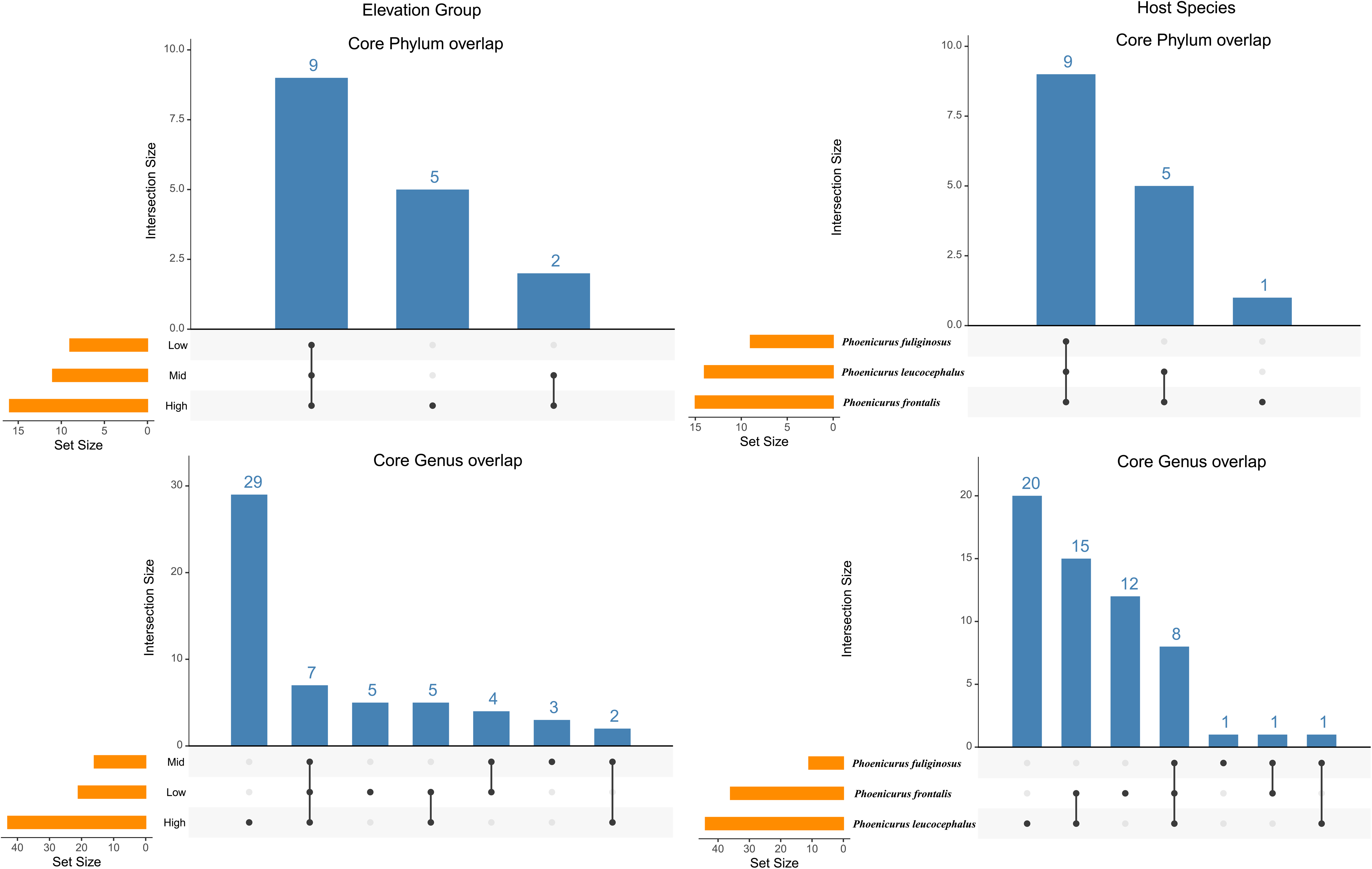
Shared and unique core gut microbial taxa across elevational groups and host species in Himalayan redstarts. UpSet plots illustrating the overlap of core bacterial phyla (upper panels) and genera (lower panels) among elevational groups (left panels) and host species (right panels). Core taxa were defined as those occurring in ≥70% of samples within a group and having a mean relative abundance of ≥0.1%. Orange horizontal bars indicate the total number of core taxa within each group (set size), whereas blue vertical bars represent the number of taxa shared by specific combinations of groups or unique to individual groups (intersection size). Connected black dots below the vertical bars indicate the groups contributing to each intersection.

**Table 2.** Number of core phyla and genera identified across host species and elevation groups. Core taxa were defined as taxa present in ≥70% of samples within a group and having a mean relative abundance of ≥0.1%.

| **Category** | **Group** | **Core phyla (n)** | **Core genera (n)** |
|---|---|---|---|
| Host species | *P. leucocephalus* | 14 | 44 |
|  | *P. frontalis* | 15 | 36 |
|  | *P. fuliginosus* | 9 | 11 |
| Elevation | High | 16 | 43 |
|  | Low | 9 | 21 |
|  | Mid | 11 | 16 |

### Species-specific and elevation-specific core genera

Analysis of host-species-specific core taxa identified 12 core taxonomic assignments unique to *P. frontalis*, including nine classified genera: *[Eubacterium] coprostanoligenes* group, *Anaerolinea, Azospirillum, Clostridium sensu stricto 1, Escherichia-Shigella, Halomonas, Mycobacterium, Proteus, and Ruegeria*. The remaining three assignments were taxonomically unresolved at the genus level **(Table S7)**. In contrast, *P. fuliginosus* had only one host-species-specific core genus, *Rickettsiella*. The largest number of host-species-specific core assignments was identified in *P. leucocephalus* (20), including nine classified genera: *Bradyrhizobium, Christensenellaceae R-7 group, Desulfovibrio, Dyella, Geothrix, Kocuria, Lactobacillus, Staphylococcus, and Tumebacillus*. The remaining 11 assignments were unclassified or incompletely resolved at the genus level **(Table S7)**. Across elevational groups, the high-elevation group contained 14 unique core genera, including *Lactobacillus*. The mid-elevation group contained three unique core taxonomic assignments: *1174-901-12*, *Methylobacterium-Methylorubrum*, and *Vibrionimonas*. The low-elevation group contained five unique core genera **(Table S6)**

### Altitude-specific microbial networks and putative keystone taxa

Genus-level microbial co-occurrence network analysis identified 28 genera connected by 35 significant positive associations in the overall network. The largest connected component comprised 20 genera and 30 associations. Based on the upper 10th percentile (≥90th percentile) of both degree and betweenness centrality, *Catenibacterium* was identified as the sole putative keystone genus in the overall network (degree = 7; betweenness = 93). Elevation-specific network analyses identified different putative keystone genera across the three elevational groups **(Fig. 7)**. *Halothiobacillus* was identified as the putative keystone genus in the low-elevation network (degree = 9; betweenness = 203), whereas *Geobacillus* was identified in the mid-elevation network (degree = 6; betweenness = 18). In the high-elevation network, *Catenibacterium* was identified as the putative keystone genus (degree = 8; betweenness = 185).

**Figure 7.**
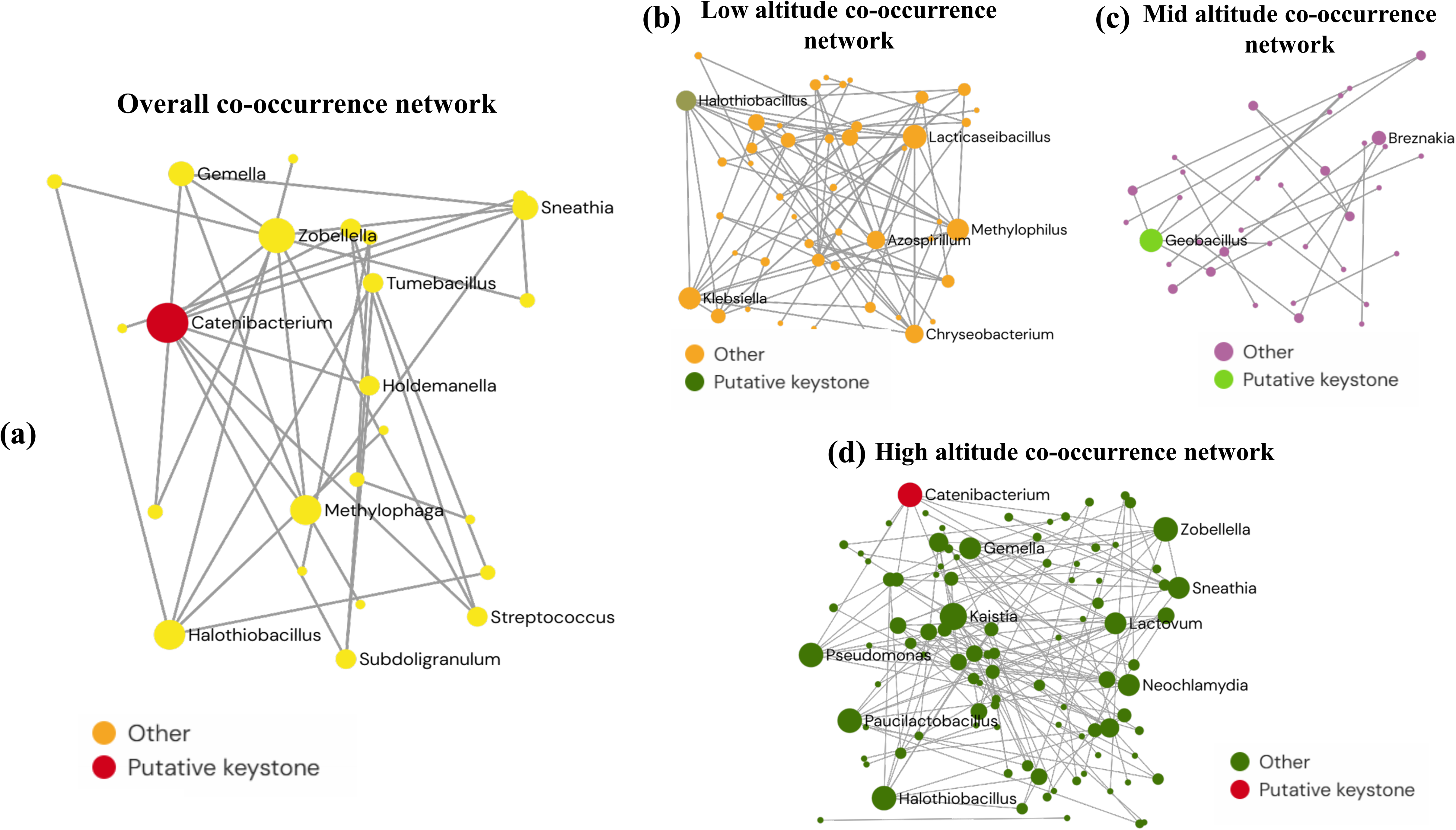
Gut microbial co-occurrence networks across elevational groups in Himalayan redstarts. Genus-level microbial co-occurrence networks constructed for the overall dataset and separately for low-, mid-, and high-elevation groups. Each node represents a bacterial genus, with node size proportional to degree centrality (number of significant associations). Networks included genera detected in ≥20% of samples within the respective dataset, with edges representing significant Spearman correlations (|ρ| ≥ 0.70, Benjamini–Hochberg-adjusted P < 0.05). Putative keystone taxa were defined as genera ranking within the top 10% for both degree and betweenness centrality in the largest connected component of each network. (a) Overall network, with *Catenibacterium* (red node) identified as the putative keystone genus. (b) Low-elevation network, with *Halothiobacillus* (green node) identified as the putative keystone genus. (c) Mid-elevation network, with *Geobacillus* (green node) identified as the putative keystone genus. (d) High-elevation network, with *Catenibacterium* (red node) identified as the putative keystone genus.

### Predicted functional profile of the gut microbiome

PICRUSt2-based functional predictions identified 28 KEGG Level 3 pathways whose predicted relative abundances differed significantly among low-, mid-, and high-elevation groups following Benjamini–Hochberg correction for multiple testing (q < 0.05; **Fig. 8; Data Table 2**). These pathways represented several functional categories, including bile acid biosynthesis, carbohydrate metabolism, amino acid biosynthesis, nucleotide and cofactor metabolism, lipid metabolism, glycan metabolism, and secondary metabolite biosynthesis.

**Figure 8.**
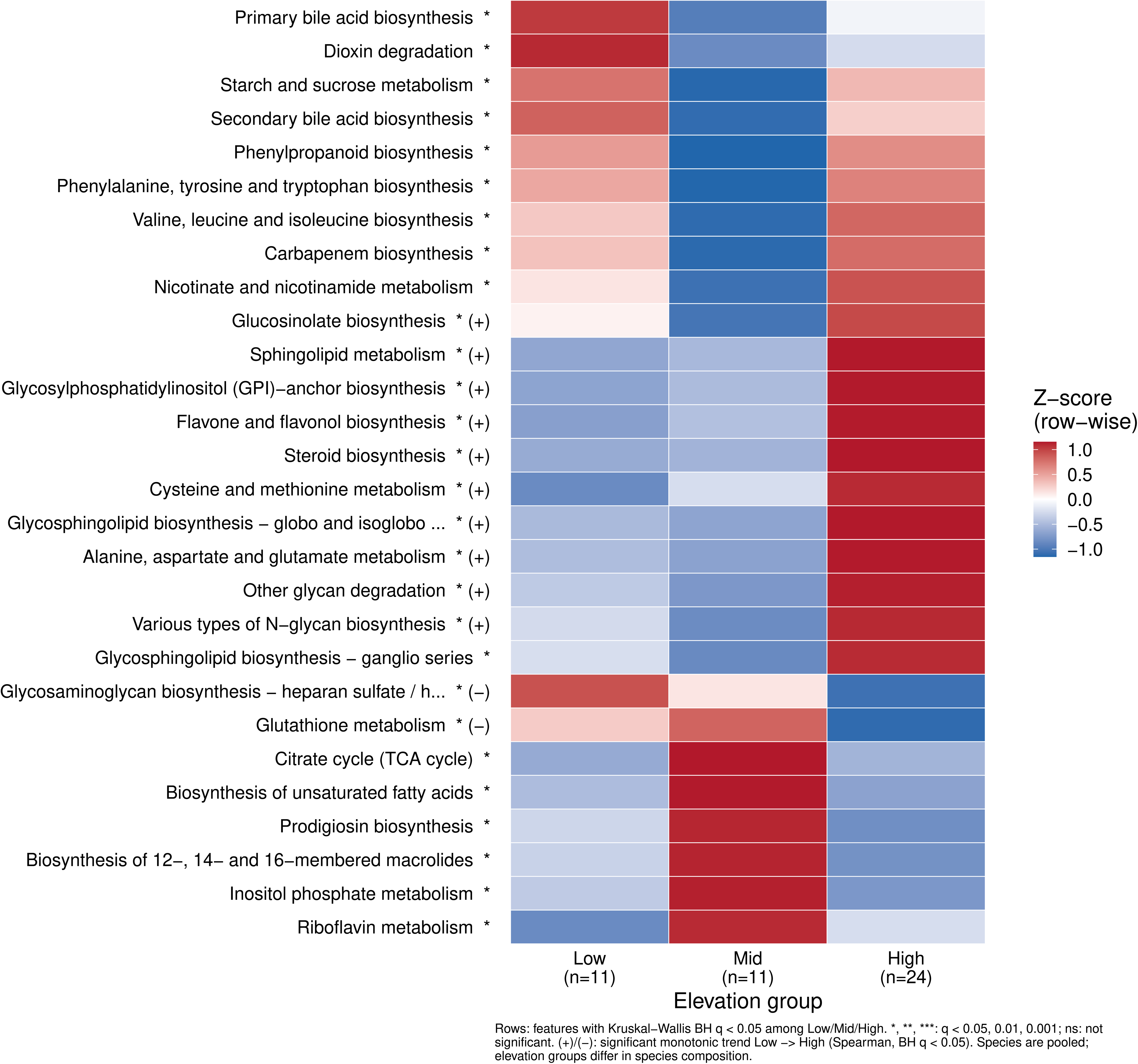
Elevational variation in the predicted functional potential of gut microbial communities in Himalayan redstarts. Heatmap showing 28 PICRUSt2-predicted Kyoto Encyclopedia of Genes and Genomes (KEGG) Level 3 pathways that differed significantly among low (n = 11), mid (n = 11), and high (n = 24) elevation groups (Kruskal–Wallis test, Benjamini–Hochberg [BH]-adjusted q < 0.05). Colors represent row-wise Z-scores of mean predicted relative abundances, with red and blue indicating relatively higher and lower values, respectively. Asterisks (*) indicate pathways with significant differences among elevational groups, whereas (+) and (−) indicate significant positive and negative monotonic associations with increasing elevation, respectively (Spearman’s rank correlation, BH-adjusted q < 0.05). Samples from all three host species were pooled within their respective elevational groups.

Spearman rank correlation analyses identified ten predicted pathways with positive associations with increasing elevation: glucosinolate biosynthesis, sphingolipid metabolism, glycosylphosphatidylinositol (GPI)-anchor biosynthesis, flavone and flavonol biosynthesis, steroid biosynthesis, cysteine and methionine metabolism, glycosphingolipid biosynthesis, alanine, aspartate and glutamate metabolism, other glycan degradation, and N-glycan biosynthesis. In contrast, glycosaminoglycan biosynthesis and glutathione metabolism showed negative associations with elevation. These associations were statistically significant in the reported analyses. Several other predicted pathways differed significantly among elevational groups without exhibiting consistent monotonic trends. These included primary and secondary bile acid biosynthesis, dioxin degradation, starch and sucrose metabolism, and riboflavin metabolism, with their highest predicted relative abundances occurring at different elevations (**Fig. 8; Data Table 2)**. Comparisons among host species did not identify statistically significant differences in predicted functional profiles after multiple-testing correction **(Data Table 3)**.

## Discussion

### Elevational variation in gut microbial diversity and community composition

Understanding how environmental gradients shape host-associated microbial communities is particularly important in montane ecosystems, where changes in temperature, oxygen availability, vegetation, and resource availability occur over relatively short geographic distances (Wang *et al*., 2022). Himalayan redstarts provide an opportunity to investigate these relationships because closely related species occupy overlapping but distinct elevational ranges and exhibit different movement strategies. In this study, we examined gut microbial communities of three Himalayan redstart species across an elevational gradient spanning 1,300–4,200 m. Elevation was significantly associated with gut microbial community composition across a gradient spanning 1,300–4,200 m, whereas host species was not a significant predictor after accounting for elevation. However, elevation explained only approximately 3% of the variation in both Bray–Curtis and Jaccard dissimilarities, suggesting that its contribution to overall microbial community structure was relatively modest. The consistency across these complementary metrics indicates that elevational variation was associated with both microbial relative abundances and taxonomic occurrence patterns. Previous studies have similarly documented elevational differences in the gut microbiomes of wild birds and mammals, although the relative importance of environmental conditions and host identity varies among systems (Suzuki *et al*., 2019; Youngblut *et al*., 2019; Bo *et al*., 2022). Environmental influences may operate through differences in diet, microbial exposure, and host physiology (Woodhams *et al*., 2020), although the present sampling design cannot distinguish the independent effects of elevation and geographic location.

In contrast to microbial community composition, alpha diversity showed no consistent elevational trend within individual host species. Although all six alpha-diversity metrics differed significantly among the combined species–elevation groups, these differences were largely associated with comparatively low diversity in *P. fuliginosus* at 2,600 m. Previous studies have reported both increasing and decreasing gut microbial diversity with elevation, potentially reflecting differences in host ecology, dietary resources, and environmental conditions (Li *et al*., 2019; Liu *et al*., 2022; Peng *et al*., 2025). Our findings suggest that elevational variation in gut microbial communities may be expressed more strongly through changes in taxonomic composition than through consistent changes in microbial richness or diversity.

### Species-specific microbial turnover and migratory ecology

Elevational patterns of microbial community composition differed between the two better-sampled redstart species. In *P. fuliginosus*, five of six pairwise elevation comparisons revealed significant compositional differences, whereas none of the pairwise comparisons within *P. leucocephalus* remained significant after multiple-testing correction. However, these differences partly reflect their distinct sampling ranges. *P. fuliginosus* was sampled from 1,300 to 3,500 m, and all significant comparisons involved at least one of the two lowest-elevation sites, whereas *P. leucocephalus* was sampled from 2,600 to 4,200 m. Distance– decay analyses revealed a contrasting pattern: microbial dissimilarity increased significantly with elevational separation in *P. leucocephalus*, but not in *P. fuliginosus*. These analyses capture different aspects of microbial variation, with pairwise community comparisons identifying differences among sampling groups and distance–decay relationships assessing whether dissimilarity increases with elevational separation. Similar distance–decay patterns have been reported in wild pikas (Li *et al*., 2019). Together, these findings suggest that microbial community turnover along elevational gradients may differ among closely related host species, although geographic separation, sampling structure, and local environmental conditions may also contribute to these patterns.

Differences in seasonal movement strategies could further influence microbial community assembly by altering exposure to environmental microorganisms and dietary resources. Migration-related changes in gut microbial communities have been documented in several bird species, including migratory warblers and thrushes (Song *et al*., 2020; Bodawatta *et al*., 2022; Schmiedová *et al*., 2023; Skeen *et al*., 2023). In our study, host species was not significantly associated with overall microbial community composition after accounting for elevation, and comparisons among species sampled within shared elevational bands revealed no significant differences in community centroids. Nevertheless, these results do not exclude an influence of migratory behavior, because movement strategies were associated with host identity and individuals were sampled during a single seasonal period. Longitudinal sampling across breeding, migration, and wintering periods would be necessary to determine whether seasonal movements contribute independently to microbial community turnover.

### Taxonomic composition, core microbiomes, and microbial co-occurrence networks

Gut microbial communities were dominated by Proteobacteria and Firmicutes, consistent with their widespread occurrence in wild birds (Grond *et al*., 2018). Their relative abundances varied among elevational groups, although few bacterial genera showed consistent elevational associations after multiple-testing correction. Notably, Lactococcus exhibited a significant negative correlation with elevation in P. fuliginosus. The relatively high abundance of Rickettsiella at mid-elevation sites may reflect dietary or environmental microbial inputs, as members of this genus are frequently associated with arthropods (Bouchon *et al*., 2011). Core microbiome analyses revealed nine bacterial phyla shared among all three host species and across all elevational categories, suggesting broad taxonomic consistency despite differences in community composition. In contrast, genus-level core membership was more variable, with eight core genera shared among species and additional taxa uniquely meeting core criteria within particular host species or elevational groups. These patterns suggest variation in recurrent microbial taxa across host and environmental contexts, although core membership does not necessarily indicate exclusive occurrence or stable intestinal colonization.

Microbial co-occurrence networks provided additional evidence of variation in community organization across elevational groups. Halothiobacillus, Geobacillus, and Catenibacterium were identified as putative keystone genera in the low-, mid-, and high-elevation networks, respectively, with Catenibacterium also identified in the overall network. The ecological characteristics of these genera differ substantially: Halothiobacillus includes sulfur-oxidizing bacteria, Geobacillus comprises predominantly thermophilic, spore-forming bacteria, and Catenibacterium includes anaerobic bacteria associated with intestinal environments (Kelly and Wood, 2000; Nazina *et al*., 2001; Kageyama and Benno, 2000). Their contrasting network prominence may reflect differences in microbial community composition, environmental exposure, or host-associated conditions. However, correlation-based networks cannot establish direct microbial interactions, and centrality-based keystone assignments do not demonstrate functional importance (Berry and Widder, 2014; Banerjee et al., 2018). These findings therefore provide a basis for identifying potentially important microbial associations rather than definitive evidence of ecological interactions.

### Predicted functional variation across elevational gradients

Predicted microbial functional profiles also differed among elevational groups. PICRUSt2 analyses identified 28 KEGG Level 3 pathways with significantly different predicted relative abundances, including pathways associated with amino acid, carbohydrate, lipid, glycan, nucleotide, and cofactor metabolism. Several pathways, including cysteine and methionine metabolism, alanine, aspartate and glutamate metabolism, sphingolipid metabolism, and glycosphingolipid biosynthesis, showed positive associations with elevation, whereas glutathione metabolism and glycosaminoglycan biosynthesis exhibited negative associations. These patterns may reflect differences in microbial substrate utilization, dietary inputs, or community composition across elevational environments. Comparable variation in predicted microbial functional potential has been reported in other high-elevation vertebrates (Li *et al*., 2019; Bo *et al*., 2025; Zhang *et al*., 2025). However, PICRUSt2 infers functional potential from taxonomic profiles and reference genomes rather than directly measuring microbial genes or metabolic activity. Consequently, these findings should not be interpreted as evidence of functional adaptation to hypoxia or other high-elevation conditions. Shotgun metagenomics and metabolomic analyses would help determine whether the predicted functional differences correspond to changes in microbial metabolic capabilities or activity.

Some limitations, however, should be considered when interpreting these findings. Elevation was confounded with geographic sampling location, preventing separation of elevational effects from site-specific environmental influences. Unequal representation of host species across elevations, particularly the limited sample size of *P. frontalis*, also constrained comparisons of host identity and movement strategies. Furthermore, fecal 16S rRNA gene sequencing may detect transient dietary or environmental microorganisms, while predicted functional profiles and co-occurrence networks provide indirect evidence of microbial ecological processes. Future studies incorporating replicated sampling sites, seasonal monitoring, dietary characterization, and metagenomic approaches will be important for disentangling environmental and host-associated influences on gut microbial communities in Himalayan birds.

## Conclusion

Our findings demonstrate that elevation is a significant, although relatively modest, correlate of gut microbial community composition in three closely related Himalayan redstart species. Elevation-associated patterns differed between host species: microbial communities in *P. fuliginosus* showed significant compositional differences among most sampled elevations, whereas those in *P. leucocephalus* exhibited increasing dissimilarity with greater elevational separation. In contrast, host species was not significantly associated with overall microbial community composition after accounting for elevation. Elevational variation was also evident in core microbial taxa, co-occurrence network structure, and predicted functional potential, particularly in pathways associated with amino acid, lipid, glycan, and carbohydrate metabolism. Together, these findings suggest that environmental conditions associated with elevational gradients may contribute to gut microbial community assembly in Himalayan birds, although the relative influences of elevation, geographic location, and host ecology remain unresolved. Future studies incorporating replicated sampling sites, seasonal monitoring across migratory periods, and shotgun metagenomics will be important for disentangling these influences and evaluating the functional significance of microbial variation in montane environments.

## Author Contributions

Conceptualization: N.D., S.L., R.K.; Investigation: P.C.; Formal analysis: A.K., P.C., S.R.; Funding acquisition: S.L., N.D., R.K.; Writing – original draft: P.C., N.D., S.R.; Supervision: S.L., R.K., S.B.; Writing – review & editing: All authors.

## Acknowledgements

We thank the Forest Department of Arunachal Pradesh for granting permission to conduct fieldwork in Arunachal Pradesh (Permit No. CWL/Gen/996/2023/Pt-III/1064-65, dated 4 July 2025). We also acknowledge the National Biodiversity Authority (NBA) for granting permission to access biological resources for research (Application No. NBA/Tech Appl/9/INBA1202204090/22/25-26, dated 2 December 2025). We are particularly grateful to Dr Umesh Srinivasan for his invaluable guidance, support, and logistical assistance throughout the study. We sincerely thank Micah Rai, Rohit Gurung, Sameer Rai, Kabir Pradhan, Bikram Manger, Sachin Dahal, Tsering Lhamu, and the late Dinesh Munda for their valuable assistance during fieldwork. N.D. was supported by the Anusandhan National Research Foundation (ANRF) National Postdoctoral Fellowship (NPDF; Sanction Order No. PDF/2025/003223, dated 24 November 2025). P.C. and S.R. were supported by the University Grants Commission (UGC) Non-NET Fellowships. Fieldwork conducted as part of a broader research project was supported in part by the National Institutes of Health (NIH), National Institute of General Medical Sciences (NIGMS), under Award Number R15GM152940 to S.L. Fecal samples analyzed in the present study were collected during this fieldwork; the NIH award did not directly support the subsequent gut microbiome analyses.

## Data Availability Statement

The data that support the findings of this study are available in the Supporting Information of this article. Raw reads are available upon request.

