## Supplementary Materials for "Elevational variation in gut microbial community composition and predicted functional potential across three Himalayan redstart species"

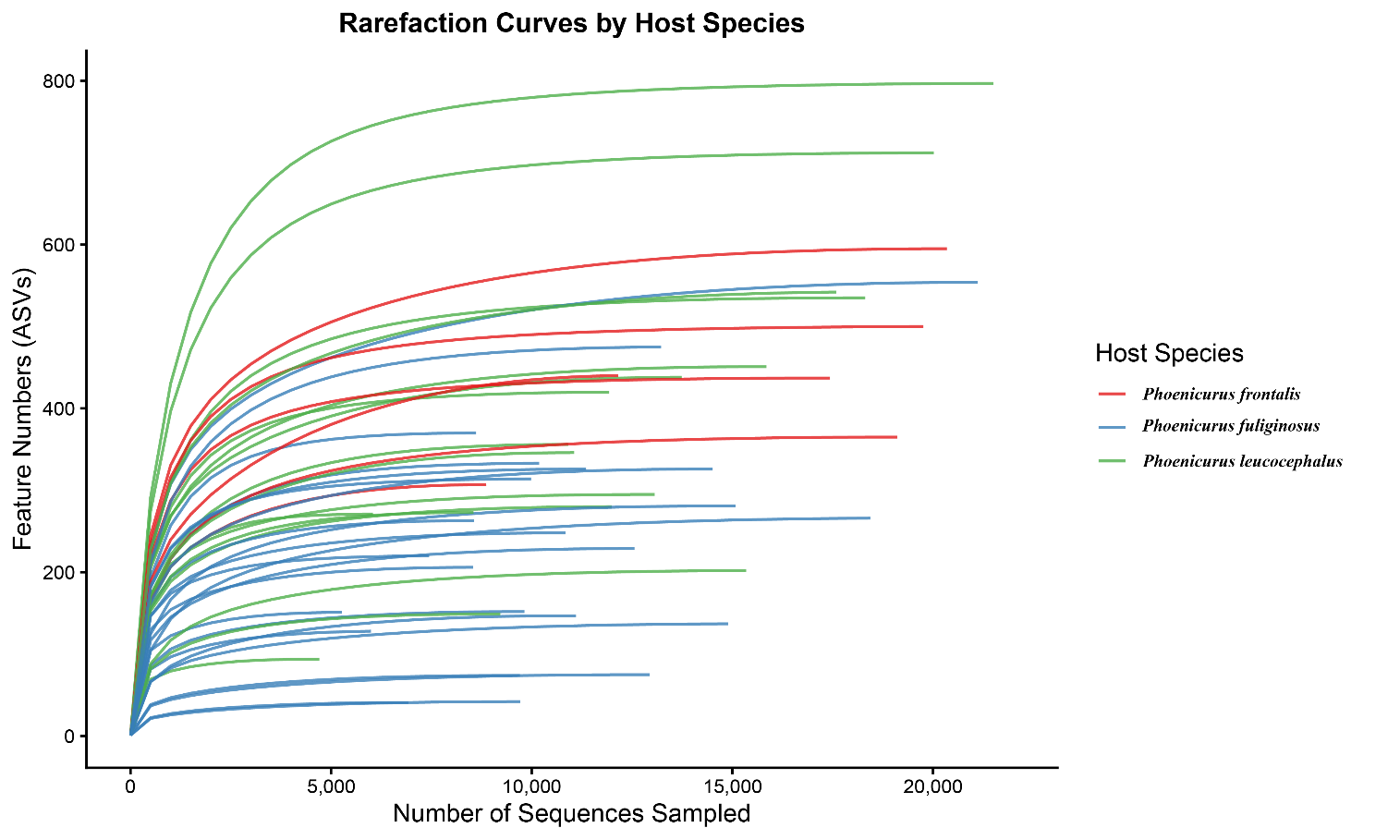

**Figure S1. Rarefaction curves of gut microbial communities.** Rarefaction curves showing ASV accumulation with increasing sequencing depth across samples. Different coloured lines represent the three redstarts species.

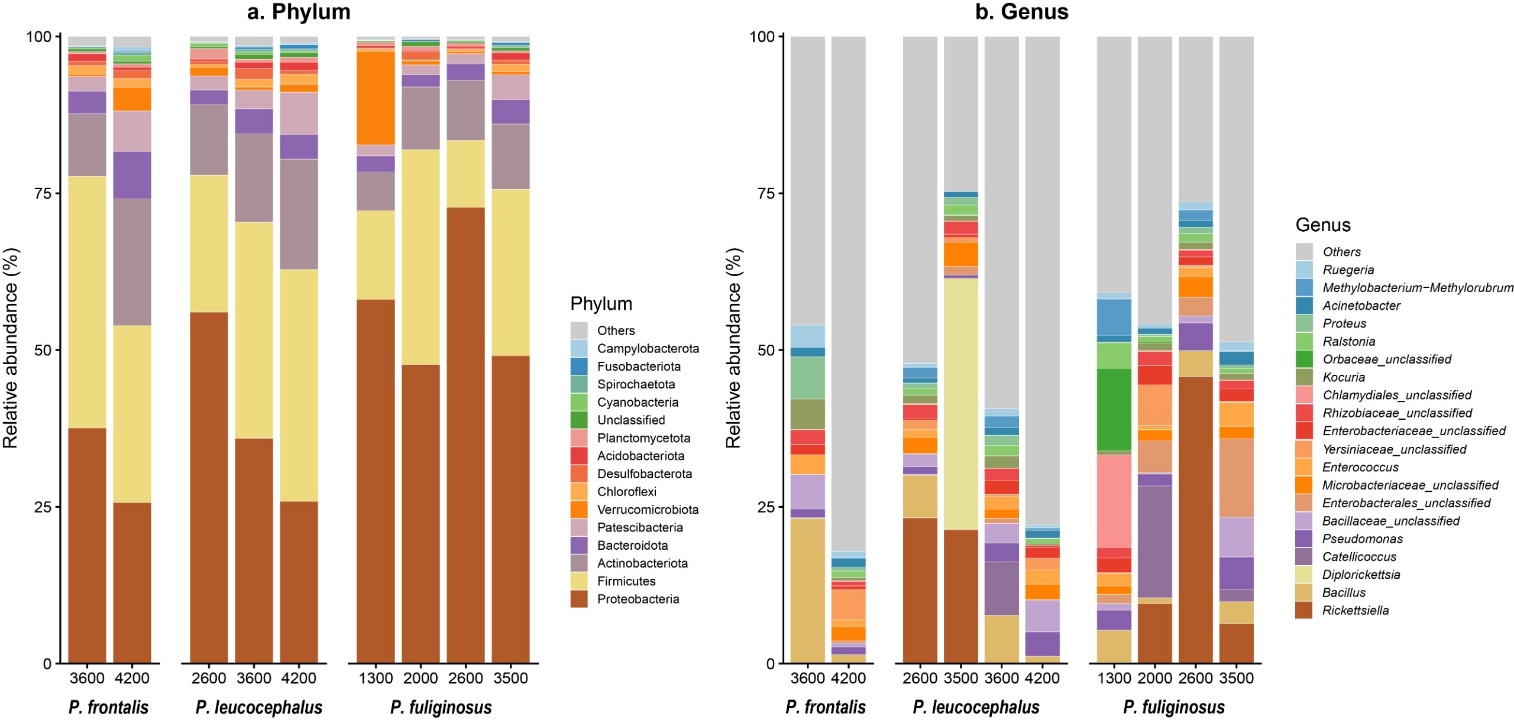

**Figure S2. Relative abundance of dominant bacterial taxa.** Relative abundance (%) of the dominant bacterial (a) phyla and (b) genera in the gut microbiome of Himalayan redstarts across the elevational gradient. Bars represent the mean relative abundance of bacterial taxa for each host species and elevation group. The 15 most abundant phyla and 20 most abundant genera are shown individually, while all remaining taxa are grouped as “Others,” representing their cumulative relative abundance.

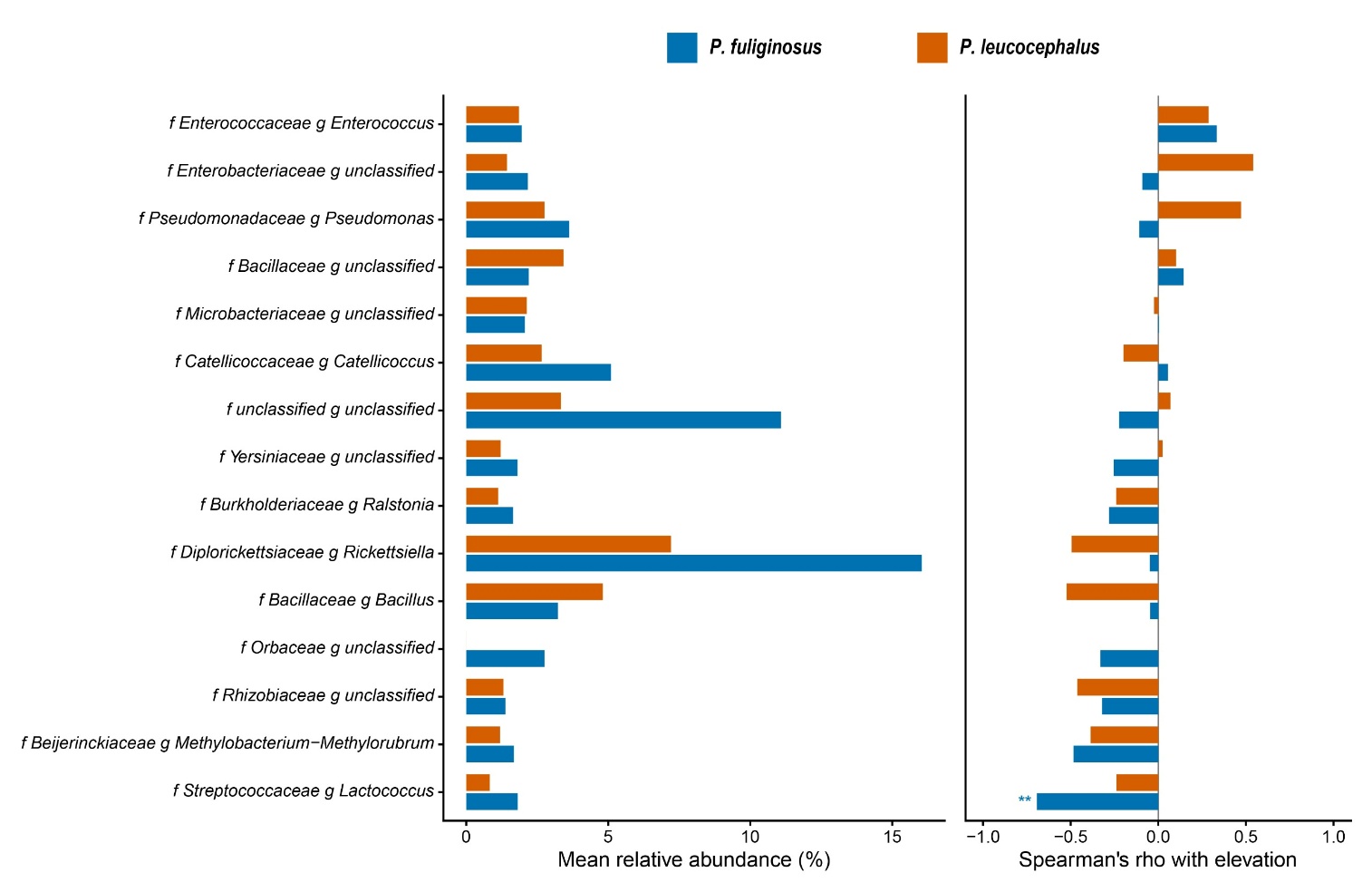
**Figure S3. Relative abundance and elevation-associated variation of dominant gut bacterial taxa.** Mean relative abundance (%) of the 15 most abundant bacterial genera in *P. fuliginosus* and *P. leucocephalus* is shown in the left panel. Genera with mean relative abundance ≥0.1% across all samples were included. The right panel shows Spearman rank correlation coefficients (ρ) between elevation and genus relative abundance for each species. Positive and negative ρ values indicate increasing and decreasing relative abundance with elevation, respectively. Asterisks indicate significant correlations after Benjamini-Hochberg correction (P < 0.01).

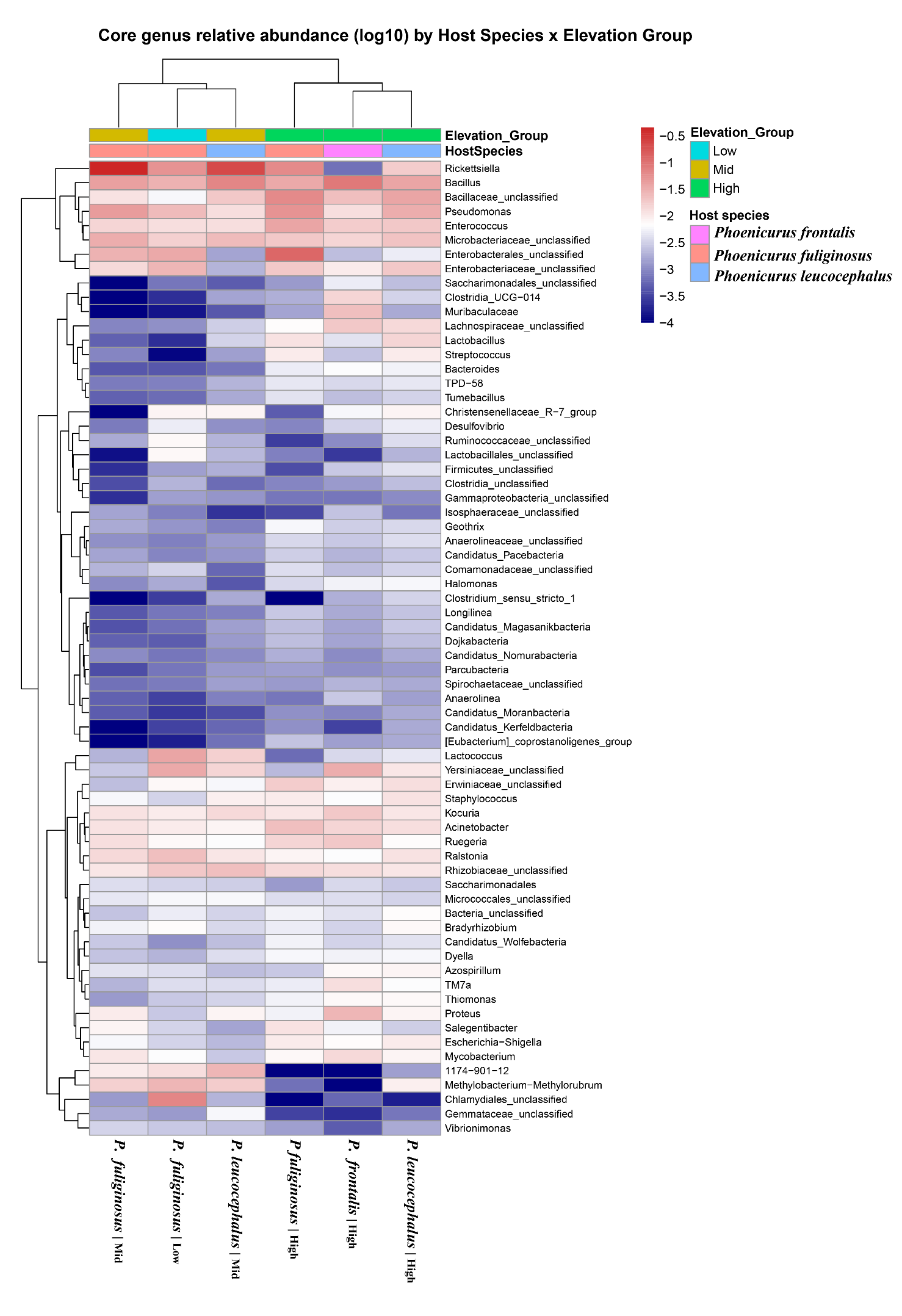

**Supplementary Figure S4. Core bacterial genus relative abundance across host species and elevation groups.** The heatmap shows the relative abundance of core bacterial genera across *P. fuliginosus*, *P. leucocephalus*, and *P. frontalis* at different elevation groups. Values are presented as log10 transformed relative abundances, with hierarchical clustering of bacterial genera and sample groups.

**Table S1: Summary of faecal samples and sampling elevations.** Summary of faecal samples and elevation groups collected for gut microbiome analysis across an elevational gradient in the West Kameng and Tawang districts of Arunachal Pradesh in August-September 2025. The host species included the resident *P. fuliginosus* and two altitudinal migrants, *P. leucocephalus* and *P. frontalis*.

| **Location Name (elevation in m)** | **Elevation Group** | **Host Species** | **Sample Sizes** |
| --- | --- | --- | --- |
| Chitre Busty - CB (1300) | Low | *P. fuliginosus* | 5 |
| Dirang - DG (2000) |  | *P. fuliginosus* | 6 |
| Jang Bridge -JB (2600) | Mid | *P. fuliginosus* | 6 |
|  |  | *P. leucocephalus* | 5 |
| Jaswantgarh - JG (3500) | High | *P. fuliginosus* | 6 |
|  |  | *P. leucocephalus* | 1 |
| Ahirgarh - AG (3600) |  | *P. leucocephalus* | 5 |
|  |  | *P. frontalis* | 2 |
| Sela Pass – SP (4200) |  | *P. frontalis* | 4 |
|  |  | *P. leucocephalus* | 6 |

**Table S2: Sequencing read processing and quality control summary of all samples.** The table presents the number of raw input reads obtained for each sample and the number of reads retained after quality filtering. It also reports the percentage of input reads that passed the filtering step, followed by the number of denoised reads and successfully merged paired-end reads. The percentage of input reads successfully merged is provided to indicate the efficiency of paired-end read merging. Finally, the number of non-chimeric reads and their percentage relative to the original input reads are presented, providing an overall summary of read retention throughout the sequence processing workflow.

| **sample-id** | **input** | **filtered** | **percentage of input passed filter** | **denoised** | **merged** | **percentage of input merged** | **non-chimeric** | **percentage of input non-chimeric** |
| --- | --- | --- | --- | --- | --- | --- | --- | --- |
| 330 | 68683 | 54555 | 79.43 | 49693 | 41936 | 61.06 | 10909 | 15.88 |
| 334 | 66033 | 51566 | 78.09 | 46588 | 38283 | 57.98 | 13735 | 20.8 |
| 336 | 35813 | 27917 | 77.95 | 25211 | 21390 | 59.73 | 6041 | 16.87 |
| 337 | 62495 | 49210 | 78.74 | 44988 | 37726 | 60.37 | 8863 | 14.18 |
| 338 | 106701 | 84909 | 79.58 | 80690 | 69535 | 65.17 | 22772 | 21.34 |
| 339 | 62677 | 48607 | 77.55 | 44685 | 37039 | 59.1 | 11927 | 19.03 |
| 340 | 75584 | 59239 | 78.38 | 53976 | 45112 | 59.68 | 15848 | 20.97 |
| 341 | 73288 | 60624 | 82.72 | 59495 | 58157 | 79.35 | 12942 | 17.66 |
| 342 | 64102 | 50874 | 79.36 | 47677 | 42098 | 65.67 | 10183 | 15.89 |
| 343 | 50334 | 40033 | 79.53 | 36595 | 30926 | 61.44 | 8611 | 17.11 |
| 345 | 87251 | 68447 | 78.45 | 64494 | 56721 | 65.01 | 18310 | 20.99 |
| 346 | 107989 | 86614 | 80.21 | 83933 | 79168 | 73.31 | 19110 | 17.7 |
| 347 | 83933 | 66217 | 78.89 | 63392 | 57596 | 68.62 | 17429 | 20.77 |
| 348 | 90076 | 70756 | 78.55 | 66568 | 56393 | 62.61 | 20017 | 22.22 |
| 349 | 94734 | 76530 | 80.78 | 71948 | 62763 | 66.25 | 13222 | 13.96 |
| 350 | 47832 | 37088 | 77.54 | 34987 | 30787 | 64.36 | 9983 | 20.87 |
| 351 | 85747 | 69079 | 80.56 | 66798 | 63254 | 73.77 | 12558 | 14.65 |
| 352 | 88840 | 72530 | 81.64 | 69906 | 64752 | 72.89 | 15915 | 17.91 |
| 353 | 23534 | 18612 | 79.09 | 17267 | 15797 | 67.12 | 4705 | 19.99 |
| 354 | 53045 | 40493 | 76.34 | 37242 | 31276 | 58.96 | 13061 | 24.62 |
| 355 | 116703 | 90591 | 77.63 | 84065 | 71270 | 61.07 | 20344 | 17.43 |
| 356 | 66132 | 48784 | 73.77 | 44816 | 39703 | 60.04 | 12153 | 18.38 |
| 357 | 50085 | 39270 | 78.41 | 36143 | 29724 | 59.35 | 11988 | 23.94 |
| 358 | 43992 | 35892 | 81.59 | 34241 | 30050 | 68.31 | 5269 | 11.98 |
| 360 | 67104 | 55160 | 82.2 | 53679 | 50316 | 74.98 | 9816 | 14.63 |
| 361 | 114208 | 93949 | 82.26 | 90456 | 84163 | 73.69 | 15072 | 13.2 |
| 362 | 148432 | 117813 | 79.37 | 111724 | 99543 | 67.06 | 21117 | 14.23 |
| 363 | 97348 | 79083 | 81.24 | 75721 | 69655 | 71.55 | 14507 | 14.9 |
| 365 | 100110 | 83044 | 82.95 | 81163 | 77815 | 77.73 | 11103 | 11.09 |
| 367 | 100495 | 75313 | 74.94 | 71662 | 62758 | 62.45 | 19758 | 19.66 |
| 369 | 117730 | 88898 | 75.51 | 83390 | 73729 | 62.63 | 17590 | 14.94 |
| 372 | 49412 | 40180 | 81.32 | 37958 | 34653 | 70.13 | 8566 | 17.34 |
| 374 | 37905 | 30368 | 80.12 | 28497 | 25119 | 66.27 | 7442 | 19.63 |
| 375 | 33470 | 27326 | 81.64 | 26690 | 26212 | 78.31 | 6923 | 20.68 |
| 376 | 73757 | 60535 | 82.07 | 58712 | 54291 | 73.61 | 5993 | 8.13 |
| 378 | 49097 | 38818 | 79.06 | 36147 | 32704 | 66.61 | 8549 | 17.41 |
| 379 | 52474 | 43259 | 82.44 | 42161 | 40769 | 77.69 | 9704 | 18.49 |
| 381 | 40481 | 32661 | 80.68 | 30594 | 27729 | 68.5 | 9215 | 22.76 |
| 382 | 38213 | 31377 | 82.11 | 30789 | 30381 | 79.5 | 9714 | 25.42 |
| 384 | 50536 | 40911 | 80.95 | 37780 | 33009 | 65.32 | 11056 | 21.88 |
| 385 | 68820 | 55167 | 80.16 | 52827 | 49036 | 71.25 | 15343 | 22.29 |
| 387 | 144188 | 119478 | 82.86 | 116573 | 111677 | 77.45 | 18437 | 12.79 |
| 388 | 50661 | 40356 | 79.66 | 38164 | 34177 | 67.46 | 8538 | 16.85 |
| 389 | 64578 | 54139 | 83.84 | 51520 | 49398 | 76.49 | 14889 | 23.06 |
| 390 | 50120 | 39534 | 78.88 | 37307 | 33006 | 65.85 | 10838 | 21.62 |
| 391 | 61369 | 48864 | 79.62 | 46330 | 41619 | 67.82 | 11352 | 18.5 |

**Table S3. Pairwise beta diversity comparisons among host species.** Gut microbial community differences were evaluated using Bray-Curtis and Jaccard dissimilarities. PERMANOVA tested differences in community composition, while PERMDISP assessed multivariate dispersion. Pairwise comparisons were performed within shared elevation bands of 2600 m, 3500-3600 m, and 4200 m. Reported statistics include PERMANOVA R² and P values and PERMDISP F statistics and P values. Significant results are indicated by P < 0.05.

| **Elevation 2600**  **Bray-Curtis** | | | | | | |
| --- | --- | --- | --- | --- | --- | --- |
| **Elevation Group** | | **PERMANOVA** | | **PERMDISP** | | |
| Group1 | Group2 | R2 | P_adj | | F | P_adj |
| *P. fuliginosus* 2600 | *P. leucocephalus* 2600 | 0.074 | 0.982 | | 0.0105 | 0.921 |
| **Jaccard**  **Elevation 2600** | | | | | | |
| **Elevation Group** | | **PERMANOVA** | | | **PERMDISP** | |
| Group1 | Group2 | R2 | P_adj | | F | P_adj |
| *P. fuliginosus* 2600 | *P. leucocephalus* 2600 | 0.099 | 0.502 | | 1.528 | 0.226 |
| **Elevation band- 3500-3600**  **Bray-Curtis** | | | | | | |
| **Elevation Group** | | **PERMANOVA** | | | **PERMDISP** | |
| Group1 | Group2 | R2 | P | | F | P |
| *P. frontalis*_3600 | *P. fuliginosus*_3500 | 0.155 | 0.55 | | 114.637 | 0.0015 |
| *P. frontalis*_3600 | *P. leucocephalus* 3600 | 0.154 | 0.559 | | 110.771 | 0.0015 |
| *P. fuliginosus* 3500 | *P. leucocephalus* 3600 | 0.077 | 0.985 | | 0.0019 | 0.974 |
| **Elevation – 3500-3600**  **Jaccard** | | | | | | |
| **Elevation Group** | | PERMANOVA |  | | PERMDISP | |
| Group1 | Group2 | R2 | P | | F | P |
| *P. frontalis* 3600 | *P. fuliginosus* 3500 | 0.151 | 0.364 | | 304.07 | 0.0015 |
| *P. frontalis* 3600 | *P. leucocephalus* 3600 | 0.151 | 0.3645 | | 462.16 | 0.0015 |
| *P. fuliginosus* 3500 | *P. leucocephalus* 3600 | 0.087 | 0.876 | | 0.163 | 0.0684 |
| **Elevation 4200**  **Bray-Curtis** | | | | | | |
| **Elevation Group** | | **PERMANOVA** | | | **PERMDISP** | |
| Group1 | Group2 | R2 | P | | F | P |
| *P. Frontalis* 4200 | *P. leucocephalus* 4200 | 0.121 | 0.052 | | 0.729 | 0.415 |
| **Elevation 4200**  **Jaccard** | | | | | | |
| **Elevation Group** | | **PERMANOVA** | | | **PERMDISP** | |
| Group1 | Group2 | **R2** | P | | F | P |
| *P. Frontalis* 4200 | *P. leucocephalus* 4200 | 0.114 | 0.116 | | 8.554 | 0.011 |

**Table S4. Relative abundance of bacterial taxa across species and elevation groups.** Relative abundance of bacterial taxa at the phylum and genus levels across three redstarts (*P. leucocephalus*, *P. fuliginosus*, and *P. frontalis*) and three elevation groups: low, mid and high elevation. Values represent the relative abundance (%) of each taxon.

| **Elevation Phylum** | |  |  |  | **Host Species Phylum** | | |  |
| --- | --- | --- | --- | --- | --- | --- | --- | --- |
| **Group** | Taxon |  | Mean_pct | SD_pct | **Species** | Taxon | Mean_pct | SD_pct |
| **Low** | Proteobacteria |  | 52.889 | 7.335 | ***P. frontalis*** | Firmicutes | 34.174 | 8.379 |
|  | Firmicutes |  | 24.187 | 14.21 |  | Proteobacteria | 31.624 | 8.423 |
|  | Actinobacteriota |  | 8.097 | 2.723 |  | Actinobacteriota | 15.107 | 7.217 |
|  | Verrucomicrobiota |  | 7.779 | 10.075 |  | Bacteroidota | 5.59 | 2.771 |
|  | Bacteroidota |  | 2.262 | 0.426 |  | Patescibacteria | 4.378 | 2.921 |
|  | Patescibacteria |  | 1.656 | 0.145 |  | Verrucomicrobiota | 2.081 | 2.434 |
|  | Desulfobacterota |  | 0.728 | 0.831 |  | Chloroflexi | 1.37 | 0.012 |
|  | Planctomycetota |  | 0.631 | 0.005 |  | Desulfobacterota | 0.984 | 0.439 |
|  | Unclassified |  | 0.483 | 0.426 |  | Acidobacteriota | 0.858 | 0.558 |
|  | Chloroflexi |  | 0.283 | 0.207 |  | Cyanobacteria | 0.469 | 0.626 |
|  | Acidobacteriota |  | 0.262 | 0.083 |  | Unclassified | 0.44 | 0.022 |
|  | Fusobacteriota |  | 0.144 | 0.179 |  | Planctomycetota | 0.439 | 0.195 |
|  | Spirochaetota |  | 0.084 | 0.102 |  | Spirochaetota | 0.402 | 0.122 |
|  | Campylobacterota |  | 0.064 | 0.056 |  | Campylobacterota | 0.393 | 0.437 |
|  | Cyanobacteria |  | 0.031 | 0.015 |  | Fusobacteriota | 0.107 | 0.099 |
|  | Others |  | 0.419 | 0.031 |  | Others | 1.584 | 0.11 |
| **Mid** | Proteobacteria |  | 64.385 | 11.824 | ***P. leucocephalus*** | Proteobacteria | 39.268 | 15.366 |
|  | Firmicutes |  | 16.262 | 7.923 |  | Firmicutes | 31.092 | 8.093 |
|  | Actinobacteriota |  | 10.427 | 1.147 |  | Actinobacteriota | 14.319 | 3.203 |
|  | Bacteroidota |  | 2.506 | 0.2 |  | Patescibacteria | 3.952 | 2.396 |
|  | Patescibacteria |  | 1.893 | 0.438 |  | Bacteroidota | 3.447 | 0.938 |
|  | Planctomycetota |  | 1.074 | 0.834 |  | Chloroflexi | 1.076 | 0.604 |
|  | Verrucomicrobiota |  | 0.889 | 0.737 |  | Verrucomicrobiota | 1.071 | 0.49 |
|  | Desulfobacterota |  | 0.364 | 0.228 |  | Desulfobacterota | 0.956 | 0.655 |
|  | Chloroflexi |  | 0.362 | 0.074 |  | Planctomycetota | 0.954 | 0.632 |
|  | Cyanobacteria |  | 0.355 | 0.263 |  | Acidobacteriota | 0.902 | 0.484 |
|  | Acidobacteriota |  | 0.354 | 0.013 |  | Unclassified | 0.607 | 0.256 |
|  | Unclassified |  | 0.28 | 0.045 |  | Cyanobacteria | 0.395 | 0.147 |
|  | Spirochaetota |  | 0.115 | 0.039 |  | Fusobacteriota | 0.356 | 0.319 |
|  | Campylobacterota |  | 0.041 | 0.021 |  | Spirochaetota | 0.299 | 0.139 |
|  | Fusobacteriota |  | 0.016 | 0.015 |  | Campylobacterota | 0.186 | 0.112 |
|  | Others |  | 0.677 | 0.249 |  | Others | 1.12 | 0.303 |
| **High** | Proteobacteria |  | 34.827 | 9.714 | *P. fuliginosus* | Proteobacteria | 56.908 | 11.515 |
|  | Firmicutes |  | 33.255 | 5.757 |  | Firmicutes | 21.387 | 10.939 |
|  | Actinobacteriota |  | 14.468 | 4.46 |  | Actinobacteriota | 9.054 | 1.949 |
|  | Bacteroidota |  | 4.615 | 1.647 |  | Verrucomicrobiota | 4.109 | 7.197 |
|  | Patescibacteria |  | 4.468 | 1.999 |  | Bacteroidota | 2.773 | 0.824 |
|  | Chloroflexi |  | 1.34 | 0.174 |  | Patescibacteria | 2.207 | 1.154 |
|  | Verrucomicrobiota |  | 1.295 | 1.449 |  | Desulfobacterota | 0.586 | 0.544 |
|  | Acidobacteriota |  | 1.049 | 0.342 |  | Acidobacteriota | 0.514 | 0.453 |
|  | Desulfobacterota |  | 0.999 | 0.482 |  | Planctomycetota | 0.507 | 0.165 |
|  | Unclassified |  | 0.584 | 0.16 |  | Chloroflexi | 0.506 | 0.444 |
|  | Planctomycetota |  | 0.472 | 0.196 |  | Unclassified | 0.436 | 0.277 |
|  | Spirochaetota |  | 0.368 | 0.081 |  | Fusobacteriota | 0.176 | 0.198 |
|  | Cyanobacteria |  | 0.336 | 0.352 |  | Spirochaetota | 0.135 | 0.114 |
|  | Fusobacteriota |  | 0.333 | 0.239 |  | Cyanobacteria | 0.083 | 0.066 |
|  | Campylobacterota |  | 0.287 | 0.243 |  | Campylobacterota | 0.075 | 0.06 |
|  | Others |  | 1.303 | 0.341 |  | Others | 0.545 | 0.202 |
|  | **Elevation Genus** | | | | **Host Species Genus** | | | |
| **Group** | Taxon |  | Mean_pct | SD_pct | **Species** | Taxon | Mean_pct | SD_pct |
| **Low** | Catellicoccus |  | 8.915 | 12.608 | ***P. frontalis*** | Bacillus | 12.234 | 15.401 |
|  | Chlamydiales_unclassified |  | 7.434 | 10.394 |  | Proteus | 3.677 | 4.361 |
|  | Orbaceae_unclassified |  | 6.62 | 9.228 |  | Bacillaceae_unclassified | 3.064 | 3.435 |
|  | Rickettsiella |  | 4.856 | 6.654 |  | Kocuria | 2.709 | 3.041 |
|  | Yersiniaceae_unclassified |  | 3.282 | 4.498 |  | Yersiniaceae_unclassified | 2.401 | 3.395 |
|  | Enterobacterales_unclassified |  | 3.275 | 2.561 |  | Ruegeria | 2.312 | 1.736 |
|  | Bacillus |  | 3.023 | 2.995 |  | Enterococcus | 2.1 | 1.397 |
|  | Methylobacterium-Methylorubrum |  | 2.964 | 3.946 |  | Rhizobiaceae_unclassified | 1.539 | 1.121 |
|  | Enterobacteriaceae_unclassified |  | 2.77 | 0.535 |  | Acinetobacter | 1.5 | 0.018 |
|  | Pseudomonas |  | 2.579 | 0.934 |  | Pseudomonas | 1.331 | 0.143 |
|  | Ralstonia |  | 2.518 | 2.203 |  | Enterobacteriaceae_unclassified | 1.145 | 0.77 |
|  | Rhizobiaceae_unclassified |  | 1.921 | 0.438 |  | Microbacteriaceae_unclassified | 1.129 | 1.597 |
|  | Microbacteriaceae_unclassified |  | 1.566 | 0.269 |  | Ralstonia | 0.487 | 0.689 |
|  | Enterococcus |  | 1.319 | 0.931 |  | Enterobacterales_unclassified | 0.172 | 0.205 |
|  | Acinetobacter |  | 1.019 | 0.098 |  | Rickettsiella | 0.052 | 0.026 |
|  | Kocuria |  | 0.938 | 0.415 |  | Catellicoccus | 0.037 | 0.052 |
|  | Ruegeria |  | 0.754 | 0.396 |  | Chlamydiales_unclassified | 0.034 | 0.048 |
|  | Bacillaceae_unclassified |  | 0.634 | 0.565 |  | Orbaceae_unclassified | 0.008 | 0.012 |
|  | Proteus |  | 0.253 | 0.136 |  | Methylobacterium-Methylorubrum | 0 | 0 |
|  | Diplorickettsia |  | 0 | 0 |  | Diplorickettsia | 0 | 0 |
|  | Others |  | 43.361 | 3.575 |  | Others | 64.069 | 25.466 |
| **Mid** | Rickettsiella |  | 34.515 | 15.977 | ***P. leucocephalus*** | Rickettsiella | 11.142 | 12.862 |
|  | Bacillus |  | 5.449 | 1.924 |  | Diplorickettsia | 10.004 | 20.007 |
|  | Microbacteriaceae_unclassified |  | 2.899 | 0.53 |  | Bacillus | 3.934 | 3.866 |
|  | Pseudomonas |  | 2.84 | 2.242 |  | Microbacteriaceae_unclassified | 2.578 | 0.968 |
|  | Methylobacterium-Methylorubrum |  | 1.706 | 0.05 |  | Bacillaceae_unclassified | 2.537 | 2.104 |
|  | Rhizobiaceae_unclassified |  | 1.658 | 0.882 |  | Pseudomonas | 2.18 | 1.551 |
|  | Bacillaceae_unclassified |  | 1.561 | 0.62 |  | Catellicoccus | 2.15 | 4.235 |
|  | Enterobacterales_unclassified |  | 1.556 | 1.999 |  | Rhizobiaceae_unclassified | 1.631 | 0.908 |
|  | Enterococcus |  | 1.402 | 0.178 |  | Enterococcus | 1.39 | 1.021 |
|  | Kocuria |  | 1.272 | 0.128 |  | Ralstonia | 1.246 | 0.41 |
|  | Ralstonia |  | 1.22 | 0.188 |  | Enterobacteriaceae_unclassified | 1.196 | 1.018 |
|  | Acinetobacter |  | 0.965 | 0.223 |  | Kocuria | 1.134 | 0.73 |
|  | Ruegeria |  | 0.945 | 0.445 |  | Yersiniaceae_unclassified | 1.096 | 0.675 |
|  | Proteus |  | 0.904 | 0.102 |  | Acinetobacter | 1.068 | 0.254 |
|  | Yersiniaceae_unclassified |  | 0.864 | 0.841 |  | Methylobacterium-Methylorubrum | 0.968 | 0.91 |
|  | Enterobacteriaceae_unclassified |  | 0.786 | 0.839 |  | Proteus | 0.937 | 0.625 |
|  | Chlamydiales_unclassified |  | 0.154 | 0.039 |  | Enterobacterales_unclassified | 0.606 | 0.583 |
|  | Catellicoccus |  | 0.05 | 0.07 |  | Ruegeria | 0.571 | 0.512 |
|  | Orbaceae_unclassified |  | 0 | 0 |  | Chlamydiales_unclassified | 0.049 | 0.089 |
|  | Diplorickettsia |  | 0 | 0 |  | Orbaceae_unclassified | 0.045 | 0.09 |
|  | Others |  | 39.254 | 18.168 |  | Others | 53.539 | 22.047 |
| **High** | Diplorickettsia |  | 6.669 | 16.336 | ***P. fuliginosus*** | Rickettsiella | 15.481 | 20.596 |
|  | Bacillus |  | 6.14 | 8.747 |  | Enterobacterales_unclassified | 5.519 | 4.922 |
|  | Rickettsiella |  | 4.642 | 8.549 |  | Catellicoccus | 4.956 | 8.635 |
|  | Bacillaceae_unclassified |  | 3.438 | 2.647 |  | Chlamydiales_unclassified | 3.749 | 7.357 |
|  | Pseudomonas |  | 2.545 | 1.777 |  | Pseudomonas | 3.681 | 1.412 |
|  | Enterobacterales_unclassified |  | 2.531 | 4.937 |  | Bacillus | 3.394 | 1.801 |
|  | Enterococcus |  | 2.048 | 1.363 |  | Orbaceae_unclassified | 3.31 | 6.557 |
|  | Microbacteriaceae_unclassified |  | 2 | 1.26 |  | Enterobacteriaceae_unclassified | 2.214 | 0.748 |
|  | Proteus |  | 1.801 | 2.485 |  | Bacillaceae_unclassified | 2.185 | 2.806 |
|  | Catellicoccus |  | 1.761 | 3.395 |  | Microbacteriaceae_unclassified | 2.091 | 0.825 |
|  | Kocuria |  | 1.608 | 1.698 |  | Enterococcus | 1.992 | 1.325 |
|  | Enterobacteriaceae_unclassified |  | 1.47 | 0.748 |  | Methylobacterium-Methylorubrum | 1.913 | 2.664 |
|  | Rhizobiaceae_unclassified |  | 1.454 | 0.799 |  | Ralstonia | 1.799 | 1.535 |
|  | Acinetobacter |  | 1.449 | 0.442 |  | Yersiniaceae_unclassified | 1.759 | 3.136 |
|  | Yersiniaceae_unclassified |  | 1.321 | 1.825 |  | Rhizobiaceae_unclassified | 1.57 | 0.501 |
|  | Ruegeria |  | 1.287 | 1.228 |  | Acinetobacter | 1.348 | 0.593 |
|  | Ralstonia |  | 0.946 | 0.593 |  | Ruegeria | 1.053 | 0.421 |
|  | Methylobacterium-Methylorubrum |  | 0.364 | 0.694 |  | Kocuria | 1.029 | 0.266 |
|  | Orbaceae_unclassified |  | 0.033 | 0.072 |  | Proteus | 0.505 | 0.35 |
|  | Chlamydiales_unclassified |  | 0.013 | 0.027 |  | Diplorickettsia | 0 | 0 |
|  | Others |  | 56.479 | 21.439 |  | Others | 40.453 | 9.911 |

**Table S5. Elevation-associated variation in bacterial genera.** Spearman rank correlations between the relative abundance of bacterial genera and elevation in *Phoenicurus fuliginosus* and *P. leucocephalus*. n represents the number of samples, rho (ρ) represents the Spearman correlation coefficient, and P value indicates the significance of the association. Negative and positive ρ values indicate decreasing and increasing relative abundance with elevation, respectively.

| **Genus label** | **Host species** | **n** | **rho** | **P value** |
| --- | --- | --- | --- | --- |
| f_Bacillaceae_g_Bacillus | *P. fuliginosus* | 23 | -0.04643 | 0.892917 |
| f_Bacillaceae_g_unclassified | *P. fuliginosus* | 23 | 0.143571 | 0.855676 |
| f_Beijerinckiaceae_g_Methylobacterium-Methylorubrum | *P. fuliginosus* | 23 | -0.48355 | 0.145546 |
| f_Burkholderiaceae_g_Ralstonia | *P. fuliginosus* | 23 | -0.28072 | 0.486144 |
| f_Catellicoccaceae_g_Catellicoccus | *P. fuliginosus* | 23 | 0.054754 | 0.892917 |
| f_Diplorickettsiaceae_g_Rickettsiella | *P. fuliginosus* | 23 | -0.04767 | 0.892917 |
| f_Enterobacteriaceae_g_unclassified | *P. fuliginosus* | 23 | -0.09102 | 0.892917 |
| f_Enterococcaceae_g_Enterococcus | *P. fuliginosus* | 23 | 0.333392 | 0.405665 |
| f_Microbacteriaceae_g_unclassified | *P. fuliginosus* | 23 | 0.004656 | 0.98318 |
| f_Orbaceae_g_unclassified | *P. fuliginosus* | 23 | -0.33014 | 0.405665 |
| f_Pseudomonadaceae_g_Pseudomonas | *P. fuliginosus* | 23 | -0.10865 | 0.892917 |
| f_Rhizobiaceae_g_unclassified | *P. fuliginosus* | 23 | -0.32107 | 0.405665 |
| **f_Streptococcaceae_g_Lactococcus** | ***P. fuliginosus*** | **23** | **-0.69323** | **0.003676** |
| f_Yersiniaceae_g_unclassified | *P. fuliginosus* | 23 | -0.25433 | 0.517655 |
| f_unclassified_g_unclassified | *P. fuliginosus* | 23 | -0.22392 | 0.57068 |
| f_Bacillaceae_g_Bacillus | *P.leucocephalus* | 16 | -0.52385 | 0.200305 |
| f_Bacillaceae_g_unclassified | *P.leucocephalus* | 16 | 0.101639 | 0.901084 |
| f_Beijerinckiaceae_g_Methylobacterium-Methylorubrum | *P.leucocephalus* | 16 | -0.38614 | 0.325745 |
| f_Burkholderiaceae_g_Ralstonia | *P.leucocephalus* | 16 | -0.2401 | 0.581252 |
| f_Catellicoccaceae_g_Catellicoccus | *P.leucocephalus* | 16 | -0.19868 | 0.645024 |
| f_Diplorickettsiaceae_g_Rickettsiella | *P.leucocephalus* | 16 | -0.49446 | 0.200305 |
| f_Enterobacteriaceae_g_unclassified | *P.leucocephalus* | 16 | 0.541035 | 0.200305 |
| f_Enterococcaceae_g_Enterococcus | *P.leucocephalus* | 16 | 0.286154 | 0.565251 |
| f_Microbacteriaceae_g_unclassified | *P.leucocephalus* | 16 | -0.02435 | 0.930464 |
| f_Pseudomonadaceae_g_Pseudomonas | *P.leucocephalus* | 16 | 0.471968 | 0.200305 |
| f_Rhizobiaceae_g_unclassified | *P.leucocephalus* | 16 | -0.4621 | 0.200305 |
| f_Streptococcaceae_g_Lactococcus | *P.leucocephalus* | 16 | -0.23852 | 0.581252 |
| f_Yersiniaceae_g_unclassified | *P.leucocephalus* | 16 | 0.023738 | 0.930464 |
| f_unclassified_g_unclassified | *P.leucocephalus* | 16 | 0.068599 | 0.930464 |

**Table S6. Elevation-specific core bacterial genera.** Elevation-specific core bacterial genera identified in *Phoenicurus fuliginosus*, *P. leucocephalus*, and *P. frontalis*. The table presents the number and taxonomic identities of unique core genera detected within each host species based on the predefined core microbiome criteria.

| **Elevation group** | **No. of core phyla** | **Core genera** |
| --- | --- | --- |
| Low | **5** | *Chlamydiales_unclassified*; *Erwiniaceae_unclassified*; *Gammaproteobacteria_unclassified*; *Lactococcus*; *Rhizobiaceae_unclassified* |
| Mid | **3** | *1174-901-12*; *Methylobacterium-Methylorubrum*; *Vibrionimonas* |
| High | **29** | *Anaerolineaceae_unclassified*; *Bacillaceae_unclassified*; *Bacteroides*; *Candidatus_Magasanikbacteria*; *Candidatus_Moranbacteria*; *Candidatus_Pacebacteria*; *Candidatus_Wolfebacteria*; *Christensenellaceae_R-7_group*; *Clostridia_UCG-014*; *Clostridia_unclassified*; *Desulfovibrio*; *Dojkabacteria*; *Dyella*; *Escherichia-Shigella*; *Firmicutes_unclassified*; *Geothrix*; *Halomonas*; *Lactobacillales_unclassified*; *Lactobacillus*; *Longilinea*; *Micrococcales_unclassified*; *Muribaculaceae*; *Salegentibacter*; *Spirochaetaceae_unclassified*; *Streptococcus*; *Thiomonas*; *TM7a*; *TPD-58*; *Tumebacillus* |

**Table S7. Host species specific core bacterial genera.** Host species-specific core bacterial genera identified in *Phoenicurus fuliginosus*, *P. leucocephalus*, and *P. frontalis*. The table presents the number and taxonomic identities of unique core genera detected within each host species based on the predefined core microbiome criteria

| **Host species** | **No. of unique core genera** | **Genera** |
| --- | --- | --- |
| *P. fuliginosus* | **1** | *Rickettsiella* |
| *P. leucocephalus* | **20** | *Bradyrhizobium*; *Candidatus_Kerfeldbacteria*; *Candidatus_Moranbacteria*; *Candidatus_Nomurabacteria*; *Candidatus_Pacebacteria*; *Christensenellaceae_R-7_group*; *Clostridia_unclassified*; *Desulfovibrio*; *Dyella*; *Geothrix*; *Kocuria*; *Lactobacillales_unclassified*; *Lactobacillus*; *Microbacteriaceae_unclassified*; *Parcubacteria*; *Ruminococcaceae_unclassified*; *Staphylococcus*; *TM7a*; *Tumebacillus*; *Yersiniaceae_unclassified* |
| *P. frontalis* | **12** | *[Eubacterium]_coprostanoligenes_group*; *Anaerolinea*; *Azospirillum*; *Clostridium_sensu_stricto_1*; *Escherichia-Shigella*; *Halomonas*; *Isosphaeraceae_unclassified*; *Micrococcales_unclassified*; *Mycobacterium*; *Proteus*; *Ruegeria*; *Saccharimonadales_unclassified.* |
